# Discovering and designing multistate protein assemblies

**DOI:** 10.64898/2026.10.26.762030

**Authors:** Chelsea N. Fries, Samuel W. K. Tipps, Adam Wargacki, Quinton Dowling, Judith Ahr, Rebecca Skotheim, Connor Weidle, Justin Decarreau, Yang Hsia, Andrew J. Borst, Neil P. King

## Abstract

Evolved protein assemblies exploit multiple weak interfaces to package cargo, facilitate metabolism, and direct cell movement, often through architectures that depart from strict geometric symmetry. Designed protein assemblies can also break symmetry through conformational flexibility or strain-based defects, but have generally lacked the capacity for large-scale structural rearrangements characteristic of evolved systems. Here, we characterize the structures and biophysical properties of emergent protein assemblies and establish a strategy for designing quasisymmetric materials that switch between assembly states. Rather than relying on intrinsic flexibility, our approach programs multiple self-assembling interfaces onto rigid oligomers. Using multistate design and ensemble predictions, we generate assemblies in which protomers can adopt alternative designed interface modes, enabling identical amino acid sequences to form distinct configurations depending on assembly pathway and environmental conditions. As a proof of principle, we construct quasisymmetric capsids that transition into micron-scale 2D arrays in response to temperature. This work establishes a strategy for encoding alternative assembly geometries within a single protein sequence and provides a foundation for engineering materials with environmentally tunable properties.

## Introduction

Nature has evolved dynamic, metastable protein assemblies that support metabolism,^1–3^ cell motility,^4–6^ and the delivery of genetic material.^7,8^ These assemblies often exploit distinct protein conformations and multiple interaction interfaces to enable structural rearrangements or break geometric symmetry to form larger assemblies and architectures than those accessible through strict symmetric packing. Bacterial microcompartments (BMCs) exemplify this principle. These large, capsid-like shells encapsulate enzymes such as ribulose-1,5-bisphosphate carboxylase/oxygenase (RuBisCO) and carbonic anhydrase, increasing the efficiency of carbon fixation.^9^ Cryo-electron microscopy (cryo-EM) has revealed that BMC shell proteins can form hexamers that interact in either planar or bent geometries, enabling shell curvature and architectural variation.^10^ This assembly strategy supports compartments large enough to encapsulate multiple enzymes while incorporating selective porosity and diverse shell geometries. Although natural assemblies such as BMCs have been engineered for specific applications,^11,12^ designing analogous materials *de novo* would enable their architecture and assembly behavior to be tailored for applications ranging from immunogen design and enzyme confinement to biologics delivery and cell signaling.

Recent advances have enabled the *de novo* design of protein assemblies that break strict geometric symmetry. Two broad strategies have emerged: pseudosymmetric heterooligomers, in which distinct amino acid sequences occupy defined positions within an assembly,^13,14^ and quasisymmetric homomers, in which intrinsic flexibility enables alternative packing geometries.^15,16^ Pseudosymmetric designs provide precise control over the geometry of each component but require distinct sequences for different structural positions, complicating production and functionalization. Flexible homomers instead use identical sequences across subunits, but their assembly behavior depends on conformational and interfacial flexibility that favors particular finite structures, including closed rings and cages.^17^ Although we have previously used controlled flexibility to design quasisymmetric protein assemblies, this strategy limits the range of accessible geometries. In particular, the conformational and interfacial constraints that favor finite structures make it difficult to design assemblies that propagate indefinitely into unbounded architectures.^18^

Inspired by natural assemblies such as BMCs, we hypothesized that explicitly designed, alternative interfaces could provide a distinct route to symmetry breaking. Rather than relying on intrinsic flexibility or deliberately introduced structural defects, this strategy encodes multiple inter-protomer interfaces on a rigid oligomeric building block. If these interfaces can support alternative assembly modes while remaining sufficiently close in free energy, the same protein sequence could participate in distinct higher-order architectures depending on assembly conditions. Implementing this strategy requires the simultaneous design of multiple compatible interfaces, control over their relative energetics, and the ability to access alternative assembly states under different environmental conditions. Such an approach could expand the range of accessible protein architectures beyond finite cages to include extended, unbounded structures.

Here, we characterize the structural and biophysical properties of emergent protein assemblies to identify design principles underlying transitions between assembly states. Using these principles, we combine multistate protein design with ensemble predictions to program multiple self-assembling interfaces onto rigid, homomeric building blocks. The resulting protomers can adopt alternative interface modes, enabling identical amino acid sequences to form distinct higher-order architectures. As a proof of principle, we design quasisymmetric capsids that transition into micron-scale 2D arrays in response to temperature. These results establish an alternative strategy for programming structural plasticity into designed protein assemblies and provide a foundation for engineering protein materials with environmentally responsive architectures.

## Results

### Emergent self-assemblies arise from interface design

We recently described pseudosymmetric nanocages in which two homomeric trimers form flat sheets on the faces of high T-number icosahedra (Figure 1A).^14^ These two trimers form flat arrays matching the design model when assembled by mixing in Tris buffers, but form nanotubes when assembled via slow removal of assembly-preventing chaotropes (Figure 1B,C). These nanotubes are of ∼200 nm diameter and up to several microns in length (Figure S1). Although we initially observed nanotubes via ‘slow’ assembly, we could also template the assembly of additional proteins using pre-formed nanotube seeds (Figure S2). Upon observing multiple assembly pathways for these two trimers, we re-examined the cryo-EM structure of the sheet-forming interface in the original icosahedra. We noted that the structure deviated from the design model via a 3 ° rotation across the assembly interface (Figure 1D), which could be sufficient to curve sheets into closed nanotubes. Extending the experimental interface structure results in a closed ring approximately 200 nm in diameter, which is consistent with the observed nanotube geometry (Figure S3). We then obtained a low-resolution reconstruction of the nanotube lattice and observed hexagonal tiling of trimers resembling the 2D array with uniaxial curvature (Figure 1E).

**Figure 1:**
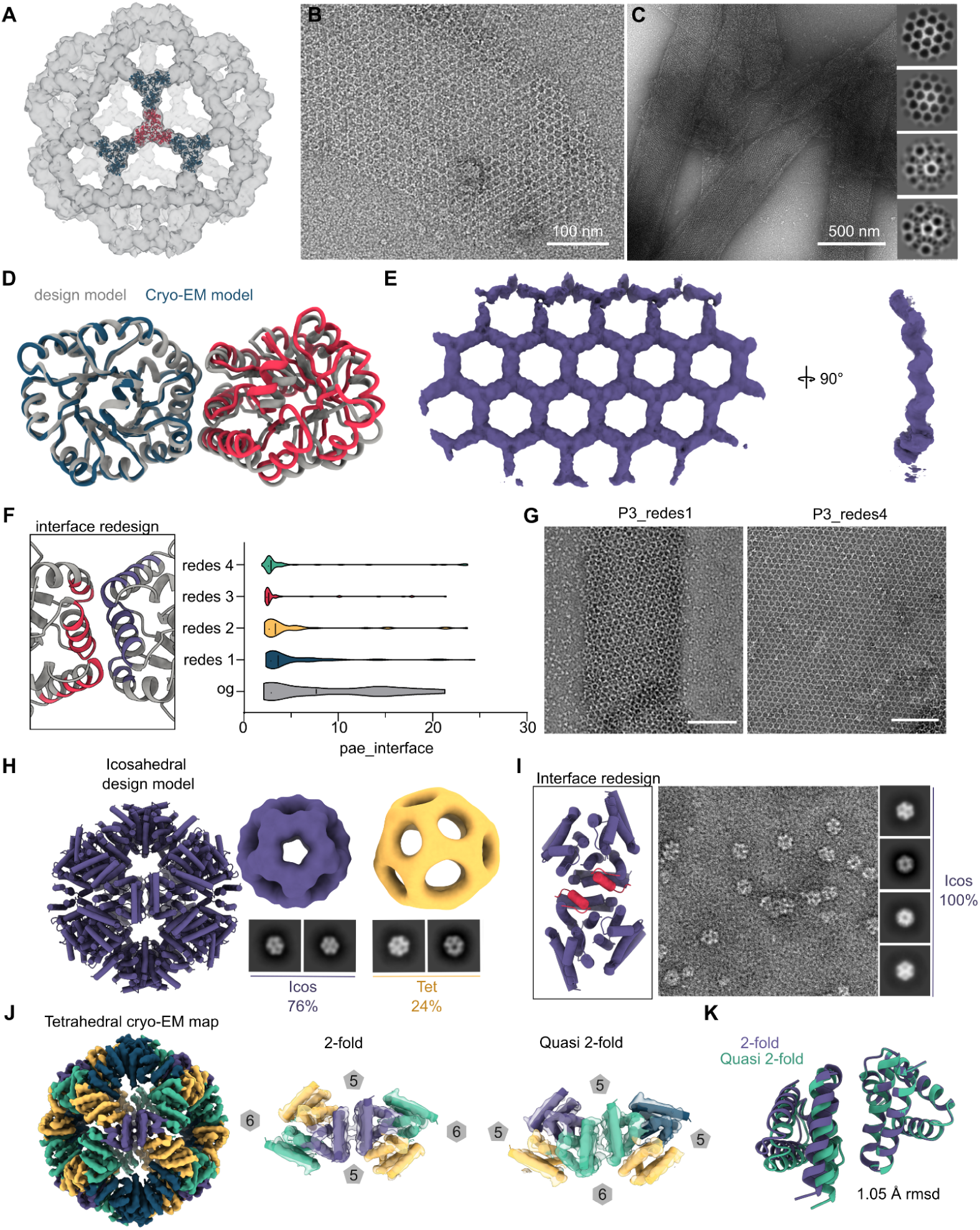
Structures and computational redesign of symmetry-breaking emergent assemblies. **(A)** Homotrimeric p3 components in I240 Goldberg polyhedra.^14^ Unique trimers are colored in blue and red, respectively. **(B)** Hexagonal 2D array formed by mixing I240 homotrimers in TBS at 20 µM. **(C)** Emergent nanotubes formed by mixing I240 trimers in TBS + 0.75% CHAPS, then dialyzing to TBS with no chaotropes. 2D class averages are shown on the right with overlapping and moiré patterning of hexagonal lattices. **(D)** Cryo-EM structure of hexon-forming interface of I240 overlaid on the symmetric p3 design model. **(E)** 9.0 Å cryo-EM map of emergent nanotubes with views of the hexagonal lattice and radial curvature. **(F)** Computational redesign of highlighted residues with proteinMPNN yields prediction ensembles with varying pae_interface distributions (n=100 predictions per sequence). **(G)** Negative stain electron microscopy (nsEM) of nanotubes (left) and 2D arrays resulting from interface redesign. Scale bar = 100 nm. **(H)** Designed icosahedral nanoparticle assembled into icosahedral and tetrahedral species as characterized by nsEM. **(I)** Interface redesign of highlighted interface helices results in single-species assemblies matching the icosahedral design model. **(J)** 4.6 Å cryo-EM map of tetrahedral species from non-optimized design with 4 quasi-equivalent positions. Two unique assembly interface structures along 2-fold and quasi 2-fold axes. The number of chains in adjacent rings is denoted in grey polygons. **(K)** Overlay of interface structures from 2-fold and quasi 2-fold positions showing a 1.1 Å RMSD deviation at the assembly interface.

To probe whether this phenomenon was related to interface design, we used ProteinMPNN to redesign residues at the trimer-trimer assembly interface and predicted the assembly with AlphaFold2 (Figure 1F). To measure the confidence of the predictions at assembly interfaces, we computed the pairwise predicted aligned error between all residues in the interface and averaged together, resulting in a metric we refer to as pae_interface. The 4 redesigned interfaces had lower pae_interface values than the original design. When we expressed these designs in *E. coli* and assembled them with slow chaotrope removal, only one of four retained the nanotube morphology, while the other 3 only formed 2D arrays (Figure 1G,S4).

In an entirely unrelated system, we observed a strikingly similar phenomenon. We generated icosahedral nanoparticles with RFDiffusion^19^ that formed both the on-target architecture and an off-target tetrahedral assembly and characterized them via nsEM (Figure 1H). Interface redesign of residues forming the 2-fold interface with ProteinMPNN resulted in monodisperse icosahedra with no observed off-target species (Figure 1I). Redesigning the interface yielded a sequence that was predicted more accurately by AF2 than the parent design (Figure S5). A 4.59 Å cryo-EM map of the tetrahedral species revealed six-membered rings and two structurally distinct assembly interfaces into which we could fit design models of individual trimers (Figure 1J). The quasisymmetric 2-fold deviated from the true 2-fold by 1.05 Å Cɑ RMSD, allowing the off-target assembly to accommodate 24 more protomers than the icosahedron (Figure 1K).

Taken together, these two systems indicated that alternative assembly geometries are related to interface design, and in some cases, differences between on-and off-target designs are reflected in AF2 prediction confidence and accuracy.

### Weak, promiscuous assembly interfaces allow protein assemblies to switch between unrelated geometries

To investigate the relationship between assembly-interface properties and alternative assembly geometries, we characterized a series of previously reported protein octahedra.^13^ The designs share the same target architecture but differ only in the residues designed at the trimer–trimer assembly interface. After expressing the designs in *E. coli* and purifying them, we obtained a structure closely matching the computational design model (Figure 2A,B). However, when we stored these assemblies at 4 °C, two designs transitioned to unexpected geometries (Figure 2C). Specifically, SL109 transitioned to form two distinct tiling patterns, one characterized by isotropic triangles and the other by alternating S-curves. Both patterns were observed in micron-scale 2D arrays, whereas nanotubes showed a regularly repeating S-curve tiling pattern (Figure S6). A second design, SL111, formed multiple fibrillar species that appear to arise from interlocked ring structures. For SL108, SL110 and SL112, no structural transitions were observed.

**Figure 2:**
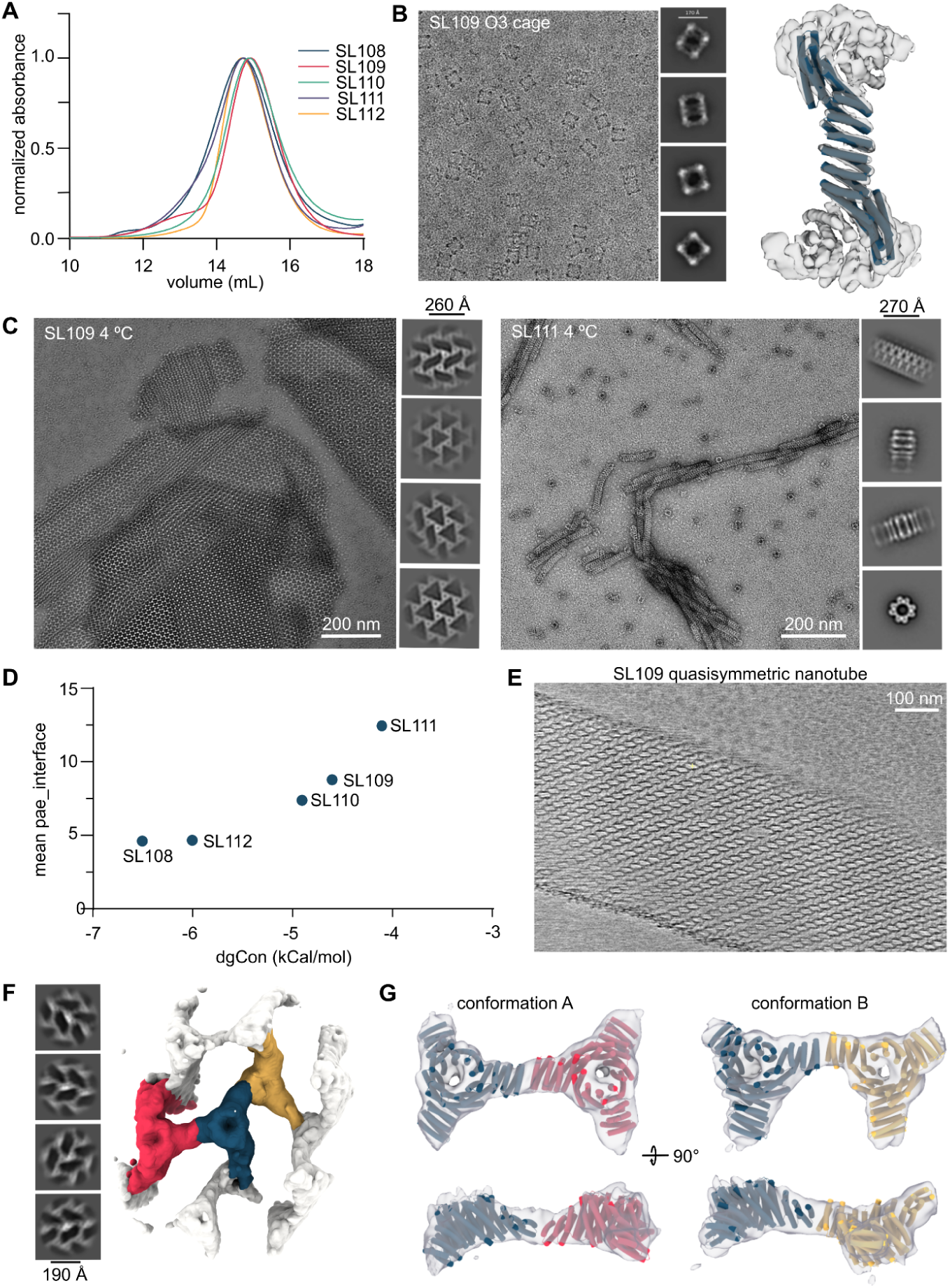
Observation and structural characterization of octahedra transitioning to unexpected geometries. **(A)** Size-exclusion chromatography (SEC) of design O3 assemblies after expression in *E. coli* and IMAC purification. **(B)** Cryo-EM micrograph, 2D classes and 3.7 Å local reconstruction with C1 symmetry of SL109 nanocage immediately after SEC purification. O3 design model fit into density using ChimeraX. **(C)** nsEM micrographs and 2D averages of SL109 2D arrays and SL111 fibrils after incubation at 4 °C. **(D)** Experimental contact energy plotted against mean pae_interface from AF2 prediction ensembles. **(E)** Cryo-ET projection of SL109 nanotubes. **(F)** 2D classes and 8.11 Å C1 cryo-EM reconstruction of SL109 nanotubes segmented to highlight individual trimers. **(G)** Trimer models docked into nanotube density show two unique lattice positions where planar trimers (blue-red) are both pointing “up” while offset (red-yellow) trimers are corrugated.

We computationally and biophysically characterized this series of designs to investigate the relationship between designed sequences and transitioning behavior. We folded each design in 5 times using models 1-5 of AF2 multimer v3 and averaged the pae_interface across the prediction ensemble. To determine if these computational differences also correlated with experimental stability, we measured the disassembly of purified cages in the presence of modest amounts of detergent (Figure S7) and calculated their contact energies as previously described.^20^ Strikingly, the contact energies obtained from experimental disassembly correlated directly with the mean pae_interface of the AF2 ensemble, where lower magnitude energy was associated with higher pae_interface (lower confidence)(Figure 2D).

The S-curved lattice formed by SL109 nanotubes showed distinctive patterning indicative of multiple types of trimer-trimer contacts. To resolve how these contacts are formed, we visualized nanotubes formed by SL109 via cryo electron tomography (cryo-ET) (Figure 2E) and obtained a 8.11 Å resolution reconstruction of the lattice using single-particle cryo-EM (Figure 2F, S8). Even at low resolution, we observed two distinct interactions within the lattice (Figure 2G). In one conformation, two trimers are coplanar with one another. In the second conformation, one trimer is rotated 180 ° relative to the other, forming a corrugated pattern of alternating upward-and downward-facing trimers.

The collection of stable and transitioning octahedra in this series further indicated that weak, low-confidence interfaces are correlated with off-target structures. In this system, weakly assembled cages reassemble into structures unrelated to the parent design. Furthermore, we observed two distinct modes of self-assembling contacts within a single architecture.

### Weakening accurate designs causes symmetry switching in protein assemblies

Because the correlation between interface strength, pae_interface, and alternative assembly geometries was striking, we asked whether weakening the assembly interface of a single-state symmetric assembly could cause it to adopt different architectures. We performed in silico alanine scans across the interfaces of SL110 and SL112, two octahedra with intermediate experimental stability (Figure 3A). We then used AF2 to predict the assembly interfaces of these mutants, obtaining a range of pae_interface values, and tested a subset of the variants experimentally.

**Figure 3:**
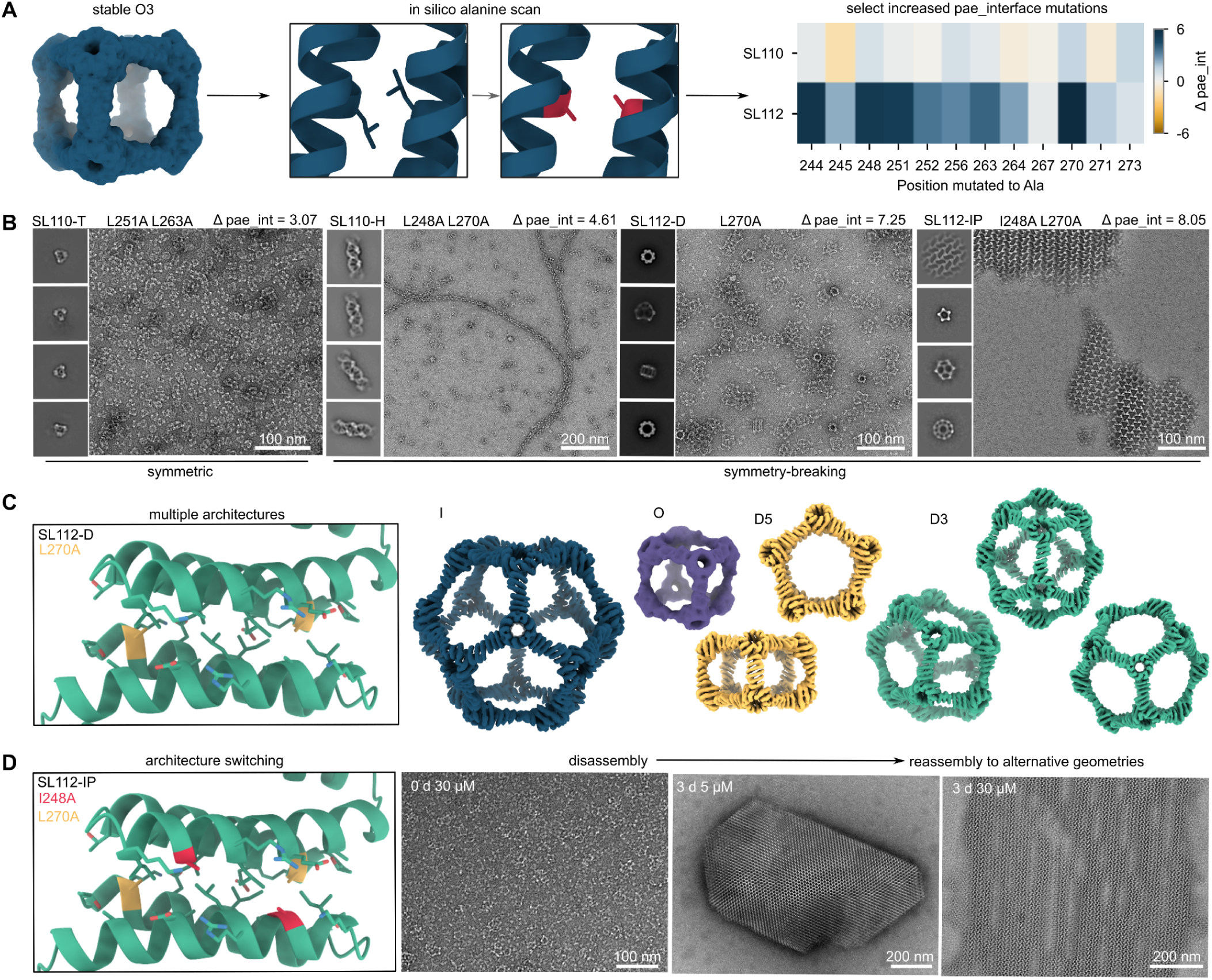
In silico alanine scan of stable octahedra produces several alternative assembly geometries. **(A)** Design procedure of weakening stable O3 assemblies SL110 and SL112 by performing an in silico alanine scan of self-assembling interfaces and predicting increases in pae_interface with AF2 ensembles. Changes in pae_interface for mutants were compared to pae_interface values for SL110 and SL112 parent designs. **(B)** nsEM of emergent assembly geometries from alanine mutants to SL110 and SL112. Mutations and Δpae_interface values are shown above micrographs and 2D class averages of each mutant. **(C)** cryo-EM maps of multiple architectures formed by SL112-D isolated in silico via heterogeneous refinement. **(D)** nsEM SL112-IP icosahedra immediately after purification and assembly into large crystalline objects when stored at 4 °C at 5 µM for 3 days and fabric-like arrays when stored at 4 °C at 30 µM for 3 days.

In several cases, we observed assembly geometries distinct from the parent octahedra (Figure 3B). Mutating two residues at the SL110 assembly interface (L251A L263A) increased the predicted pae_interface and promoted a switch from an octahedron to a tetrahedron (SL110-T). Variants with larger increases in pae_interface formed multiple species after expression in *E. coli*, some of which were quasisymmetric. Mutating residues 248 and 270 in SL110 induced formation of a double-helix structure (SL110-H, L248A L270A). A single mutation, L270A, produced multiple SL112-D species in addition to the parent O3 structure. Introducing an additional mutation at residue 248 yielded SL112-IP (I248A L270A), which formed icosahedra and 2D arrays composed of triangular and S-curve lattice structures.

We characterized several geometric species isolated from a single expression of SL112-D (Figure 3C) composed of 4-and 5-membered rings (denoted as 4^x^5^y^ assemblies to describe the number of tetrons and pentons in each assembly, respectively). We used cryo-EM to computationally isolate and reconstruct icosahedral 5^12^ cages, D3-symmetric 5^6^4^3^ cages, D5-symmetric 5^2^4^5^ cages, and octahedral 4^6^ cages (processing details in Figure S9). These assemblies exhibited multiple species and, within individual species, multiple protein conformations, with 4-and 5-membered rings coexisting within the same structure.

SL112-IP displayed switching between overall geometries in a manner akin to SL109. After expression and purification as an icosahedra, we observed condition-dependent transitions to multiple unbounded species at 4 °C (Figure 3D). We also reconstructed 3D tomograms to visualize layers of the lattices formed by SL112-IP (Figure S10) and observed twisted stacks of triangular lattices and flat lattices of S-curves similar to those observed for SL109. In this case we observed multiple species which are formed at varying times, temperatures, and concentrations, classifying this design as a multistate material.

### Alternative symmetric assemblies sample multiple interface conformations

After observing substantial changes in assembly geometry upon weakening assembly interfaces, we asked whether the new structures arose from protein-subunit flexibility, shifts in interface geometry, or both. We hypothesized that the new symmetric species would exhibit shifts in interface geometry relative to the parent O3 assemblies and that distinct positions within a single quasisymmetric assembly would differ from one another.

We obtained a cryo-EM map of SL110-T tetrahedra at 3.3 Å resolution and locally refined the trimer–trimer interface to 2.4 Å (Figure 4A, S11). We used ISOLDE^21^ to relax models of SL110 protomers into the map density, resolving an inter-protomer contact involving F244 as well as backbone movements. Alignment of the experimental structure to the SL110 parent design model revealed a Cα RMSD of 2.3 Å for the protomer and 6.8 Å for the entire complex (Figure 4B). To examine interface deviations directly, we isolated the four terminal helices from each model and aligned them. This comparison revealed translation and rotation of the terminal helices relative to the parent design model.

**Figure 4:**
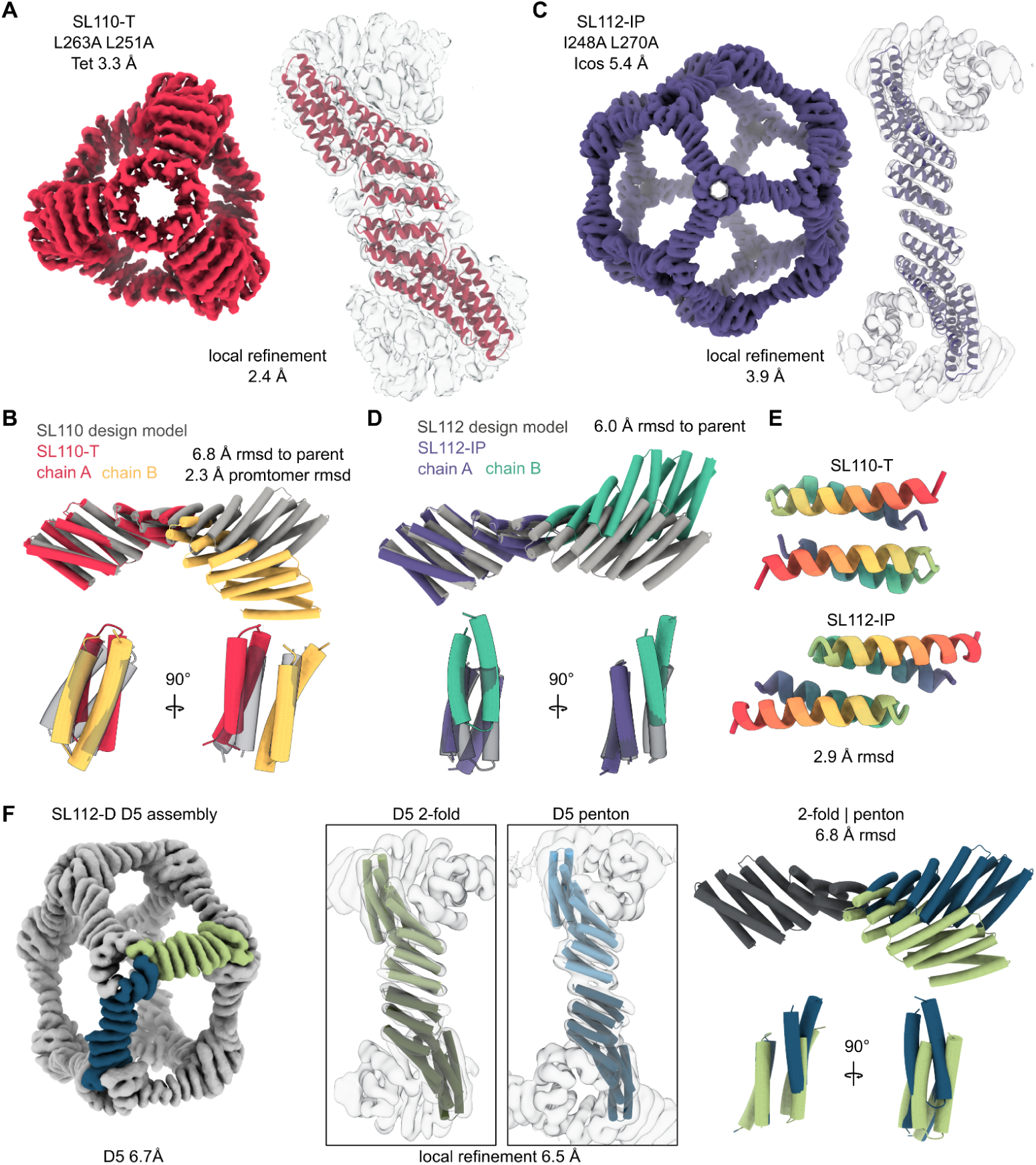
Alternative structures are a result of deviations in inter-oligomer contacts at assembly interfaces. **(A)** Cryo-EM map of SL110-T at 3.3 Å global resolution with tetrahedral symmetry. Local structure of assembly interface refined to 2.4 Å. Design model protomers were relaxed into density using ISOLDE to obtain a model of interprotomer positioning and side chain density for F244. **(B)** SL110-T assembly interface colored in yellow and red aligned to a single chain of SL110 parent interface colored in grey. The SL110-T protomer deviates from the parent design-model protomer by 2.3 Å Cα RMSD; the SL110-T complex deviates from the parent complex by 6.8 Å Cα RMSD. **(C)** Cryo-EM map of SL112-IP refined to 5.4 Å with I symmetry. Local density of assembly interface refined to 3.9 Å. Design model protomers were fitted into density using ChimeraX to obtain a model of protomer positioning. **(D)** SL112-IP assembly interface colored in green and purple aligned to a single chain of SL112 parent interface colored in grey. SL112-IP deviated from the parent complex by 6.0 Å Cα RMSD. For both B and D, two terminal helices are overlaid on one another to visualize shifts at the assembly interface. **(E)** Two terminal helices of SL110-T and SL112-IP models colored from N-terminus (red) to C-terminus (blue). **(F)** Cryo-EM map of two quasiequivalent positions in D5 assemblies formed by SL112-D refined to 6.5 Å. Design model protomers were fit into density using ChimeraX to obtain a model of protomer positioning. Penton and 2-fold symmetric contacts deviate from one another by 6.8 Å Cα RMSD.

Similarly, we determined the structure of icosahedral nanocages formed by SL112-IP after bacterial expression, before their transition to larger arrays. The overall cage map was resolved to 5.4 Å, and the trimer–trimer interface was locally refined to 3.9 Å (Figure 4C, S12). We fitted SL112 protomers into the map density and observed a relative displacement of the two protomers across the assembly interface, with a Cα RMSD of 6.0 Å compared with the parent SL112 design model (Figure 4D). Examination of the terminal helices revealed a relative slip between the two helices.

The SL110-T and SL112-IP structures exhibited distinct changes in their assembly interfaces. When aligned side by side (Figure 4B, D), SL110-T showed a downward lever-arm motion, whereas the SL112-IP chains rotated upward in the same viewing orientation. The terminal helices largely overlapped in SL110-T but were staggered in SL112-IP, which also had a smaller interface contact surface (Figure 4E).

To determine whether distinct interfaces could also occur within a single assembly, we analyzed two nonequivalent positions in the D5-symmetric 5²4⁵ cages formed by SL112-D. We obtained a cryo-EM map of the D5 assemblies at 6.7 Å resolution and locally refined the trimer–trimer contacts to 6.5 Å (Figure 4F, S13). Alignment of the two interfaces revealed a substantial shift in interface geometry accompanied by lever-arm movement of the opposing chain. The two trimer–trimer complexes differed by 6.8 Å RMSD despite occurring within the same assembly.

Across these assemblies, we observed substantial deviations in assembly-interface geometry, suggesting that interface remodeling could be a target for computational design. Notably, when we used AF2 to predict these assemblies 100 times, it did not reproduce the experimentally observed alternative structures (Figure S14). We therefore hypothesized that this structural phenomenon could be recapitulated through the design of interfaces that accommodate alternative conformations, and that sequences designed for these structures would be more likely to yield the experimentally observed architectures than sequences obtained through serendipitous mutations.

### Emergent quasisymmetric assemblies can be stabilized via computational design

We observed many symmetry-breaking and state-switching structures serendipitously, but exploiting these features requires the ability to design and predict such structures computationally. Using the geometries of emergent symmetry-breaking structures as a test case, we sought to redesign assembly interfaces to accommodate multiple conformations within the same structure.

We used backbone conformations derived from cryo-EM models to stabilize the D3-symmetric cages formed by SL112-D (Figure S15). We extracted the backbones of two chains at edge A, which forms only a 5-membered ring, and edge B, which is shared by 4-and 5-membered rings. We then used ProteinMPNN to design residues at the assembly interface of either edge while requiring the same amino acid identities at corresponding positions in both conformations (Figure 5A). We predicted 47 sequences using AF2 and selected those that predicted both state A and state B with an RMSD of less than 3 Å across models 1–5. AF2 did not predict the experimentally observed structure of the parent SL112-D sequence in any of 100 predictions (Figure S14). In contrast, when we evaluated only five AF2 predictions per designed sequence, 15 sequences were predicted to adopt both state A and state B (Figure 5B, S16).

**Figure 5:**
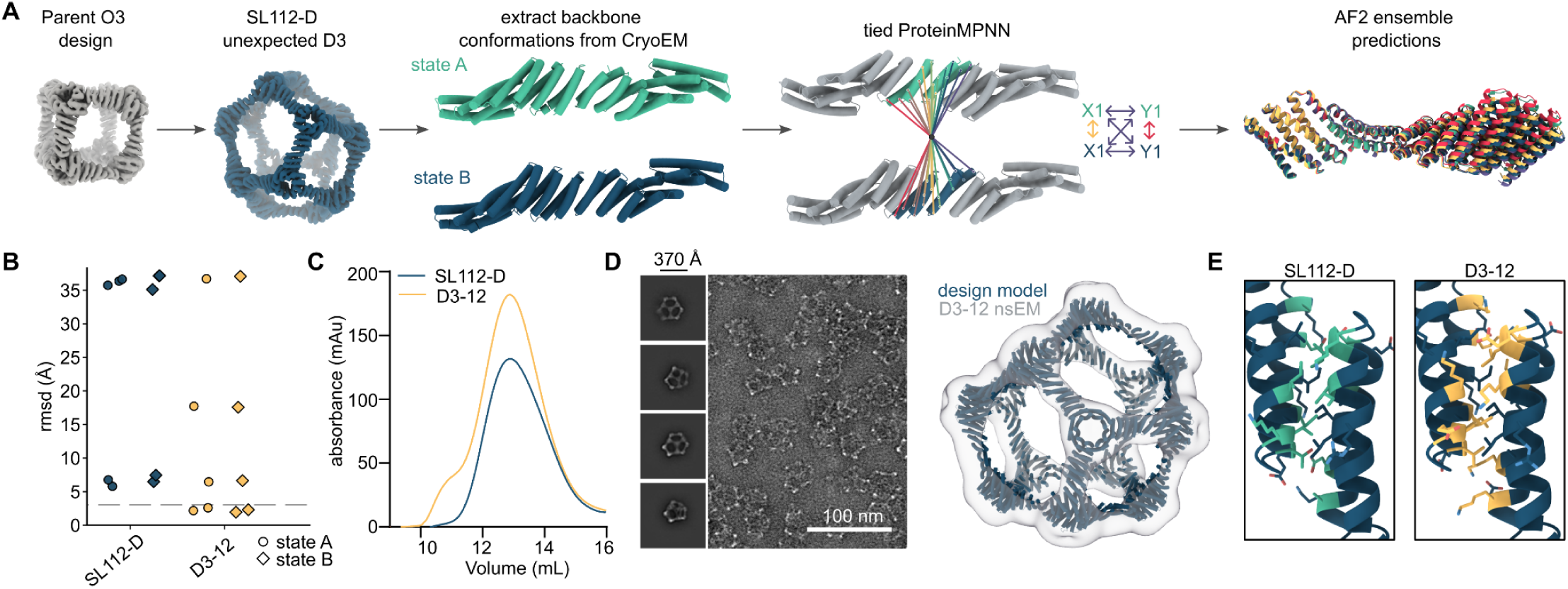
Multistate interface design generates predictable sequences for emergent quasisymmetric structures. **(A)** Design procedure to stabilize the geometry of emergent D3 nanocages formed by SL112-D. Briefly, backbone conformations were derived from the cryo-EM model, protomers of 4-5 edges and 5-5 edges were defined as state A and state B respectively. Tied ProteinMPNN was used to design residues at the same position of both chains in both states at interprotomer contacts. Sequences were then predicted with 5 models of AF2 and filtered for RMSD less than 3 Å to both state A and state B design models. **(B)** RMSD of AF2 predictions of SL112-D and D3-12 aligned to design models. (n=5 predictions per sequence, models 1-5 AF2 multimer v3) **(C)** SEC of nickel-purified SL112-D and D3-12 assemblies using an S6 column. **(D)** nsEM of D3-12 assemblies and reconstruction matching D3 design model. **(E)** Models of interface-forming residues in SL112-D emergent designs and D3-12 designed contacts.

After expression in *E. coli*, we identified designs that eluted at the same volume as SL112-D (Figure 5C, S17). We selected the best-expressing and most monodisperse design for further characterization and observed a structure matching the D3-symmetric 5⁶4³ cage design model (Figure 5D, S18). This design had a distinct sequence, with substitutions at eight positions relative to SL112-D (alignment shown in Figure 5E, S19). These results indicate that multistate interface design can generate sequences that form symmetry-breaking architectures and may be applicable to other target structures.

### Multiple interfaces can be designed simultaneously to *de novo* design quasi-symmetric assemblies

Building on our observations of emergent multistate assemblies, we sought to design assemblies with multiple states that were predicted by AF2, with low confidence across interface residues in both states. We hypothesized that mimicking the structural features of emergent state-switching assemblies would enable us to design protein assemblies capable of switching between two states and forming quasisymmetric structures. As a test case, we designed dimers and trimers (Figure 6A,B) to assemble into two target states corresponding to hexon-forming planar geometries (state A) and penton-forming geometries within an icosahedral assembly (state B) (Figure 6C). By restricting rotation of the trimeric component between these states, we aimed to enable the components to tile into extended 2D arrays or icosahedral assemblies. These geometries are compatible with idealized T=4 icosahedra (Figure 6C), but because of their quasisymmetric nature can also tile into irregular capsid geometries, much like evolved capsids.^22,23^

**Figure 6:**
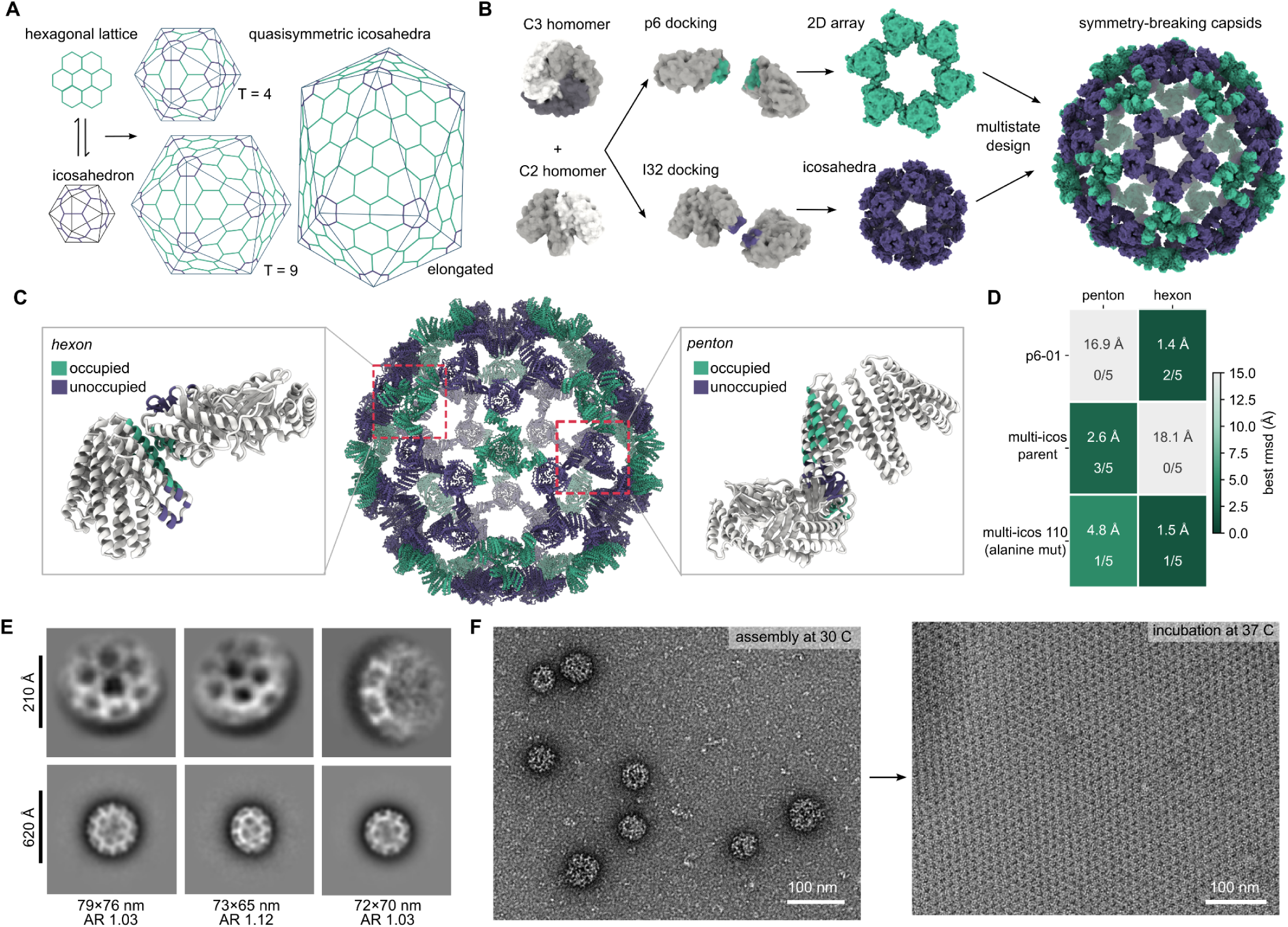
Multiple contacts can be designed *de novo* to generate dynamic symmetry-breaking materials. **(A)** Geometry of quasisymmetric capsids can occur via co-organization of hexagonal lattice and pentagonal defects. **(B)** Design strategy of separate docking and sequence design of C2 and C3 symmetric oligomers to generate orthogonal interface patches for pentons and hexons within high T-number capsids. **(C)** Design model for multi-icos 110 nanocages in a hypothetical T=4 architecture. **(D)** Heat map of best AF2 predictions for hexon and penton design states for each step of the design process with the RMSD of the best prediction and the fraction of 5 predictions under 5 Å RMSD noted in each box. **(E)** 2D class averages of lattice geometry and complete capsids with size and aspect ratio listed below each class. **(F)** Transition of nanocages assembled at 30 °C into large 2D arrays following incubation at 37 °C.

We first designed and experimentally validated a single-state 2D hexagonal lattice (state A) using an MPNN-redesigned variant of the trimer 3jv1 and a de novo helical repeat protein. We docked the two oligomers as rigid bodies using RPXDock^24^, designed interface residues with ProteinMPNN,^25^ and folded one chain from each protomer using AF2-Multimer.^26^ We selected designs whose predicted structures matched the design model with an RMSD of less than 2 Å. Of the 38 designs tested experimentally, one design named P6-01 reliably assembled into 2D arrays, with no off-target structures observed (Figure S20). With this validated hexagonal lattice as one assembly state, we next sought to design a second state by docking the same dimer and trimer into an icosahedral arrangement and designing the corresponding assembly interface with ProteinMPNN. We filtered for sequences that accurately predicted the icosahedral penton geometry and introduced targeted alanine mutations to weaken this interface. We selected pairs of residues with low PAE in designs that predicted the penton state, with the aim of weakening the icosahedral interface in silico. Following these mutations, sequences that had previously predicted only the penton state were predicted to adopt both hexon and penton conformations, identifying them as multistate interfaces in silico. Specifically, the parent sequence P6-01 predicted only the hexon, then adding an interface for the icosahedral geometry (multi-icos parent) predicted only the penton, but a single alanine mutation on each assembly component (multi-icos 110) predicted both the hexon and the penton (Figure 6D). Designs predicted to adopt both state A and state B with an RMSD below 5 Å were ordered, expressed in *E. coli*, and assembled in equimolar ratios.

We tested 63 designs and identified one that assembled into spherical objects ranging from 50 to 100 nm in size. 2D class averages showed neighboring hexons consistent with the design model, and we observed several closely related species with varying sizes and aspect ratios (Figure 6E). These classes showed regularly spaced features around the capsid perimeter, and features on the capsid face consistent with the design model could also be resolved (Figure 6F).

In addition to forming irregular capsids consistent with our quasisymmetric design scheme, these assemblies exhibited metastability, with their overall geometry depending on assembly conditions. Higher assembly concentrations and temperatures favored the formation of 2D arrays, whereas lower temperatures and concentrations favored nanocages (Figure S21). Once assembled, nanocages could also transition into large, unbounded 2D arrays when incubated overnight at 37 °C (Figure 6F).

## Discussion

Here, we establish multistate interface design as a strategy for engineering protein assemblies that break symmetry and switch between distinct geometries. By characterizing emergent assemblies, we identified quasisymmetric geometries and weak interfaces as features associated with state switching. We used these observations to design assemblies with multiple interfaces predicted by AF2 to support alternative assembly states, then applied this strategy to de novo protein assemblies that form quasisymmetric capsid-like structures and transition into unbounded 2D arrays. These results demonstrate how multiple designed interfaces can enable a single protein sequence to support alternative assembly geometries.

This approach enables protein assemblies to adopt two well-defined structural states, advancing beyond strategies that toggle between assembly and disassembly. In designed protein assemblies, pH sensitivity^27^ and conditional interfaces^28^ have been used to control assembly and disassembly, creating dynamic systems. Nature uses molecular switches to regulate protein assemblies during processes such as changes in shape, genetic material delivery, and cargo release.^29,30^ Our work represents a step toward designing materials that switch between distinct, well-defined assembly geometries.

The symmetry-breaking mechanisms observed here differ from those in previously described quasisymmetric designs.^15–17^ Whereas prior approaches have relied on subunit flexibility to accommodate alternative packing geometries, our results demonstrate that multistate interfaces provide an additional route to symmetry breaking. The relatively small deviations in protomer structure observed in SL109, P3 subunits, and our designed state-switching assemblies, compared with previously described designs (Figure S22), suggest that symmetry breaking can be achieved without substantial changes in protomer conformation. Together, these findings establish multistate interface design as a complementary strategy for engineering assemblies with alternative geometries and potentially unbounded architectures.^18^

This strategy provides a foundation for increasingly predictable design of state-switching materials. Key opportunities include defining how alternative assembly states should relate to one another and developing methods to predict the energetics and specificity of weak, switchable interfaces. Because static structures do not explicitly capture assembly dynamics and transition states are less readily defined for protein assemblies than for enzymatic reactions,^31,32^ new computational methods could further expand our ability to model these systems. Ultimately, designing transition pathways between assembly states could enable finer control over material switching and potentially allow chemical or mechanical work to be performed as subunits rearrange.

## Acknowledgements

We thank Cyrus Haas, Susan Kleinfelter, Helen Eisenach, Alexis Courbet, Shingo Honda, and Adam Broerman for helpful discussions. We thank Sangmin Lee and Ryan Kibler for sharing plasmids and insights into designed octahedra. We thank Rose Marie Yasuda for collecting cryo-EM data at the Pacific Northwest Cryo-EM Center. We thank Joel Quispe, Sasha Dickinson, Fangfang Zhang and Wentao Jiang for the management of the University of Washington Arnold and Mabel Beckman Cryo-EM Center.

## Author Contributions

CNF conceptualized the studies, synthesized and screened assemblies, collected and processed cryo-EM data, and wrote the manuscript. ST collected and processed cryo-EM data and wrote the manuscript. JA synthesized and screened protein assemblies. AW analyzed data and performed biophysical experiments. RS, CW, and AB collected and processed cryo-EM data. YH designed building blocks and contributed to conceptual discussions. JD performed experiments on nanotube templating. QD conceptualized phenomena and characterized assembly structures. NPK conceived the study, provided insight, and revised the manuscript.

## Methods

### Prediction of assembly interfaces with AF2

Assembly interfaces were predicted with AlphaFold 2 Multimer v3 using ColabFold. Slightly different settings were used for different design problems to achieve a signal-to-noise ratio relative to computational controls. For P3 assemblies and diffused icosahedra, sequences were predicted using the original design model as an “initial guess.”^26^ For multistate design, an initial guess was not used. P3 assemblies and multistate capsids were predicted using a multiple sequence alignment (MSA) because the building blocks are derivatives of native proteins. All other designs were predicted in single-sequence mode with no MSA. For SL octahedra and downstream designs, we solubilized the trimerization domain in silico to prevent the oligomerization interface from dominating the prediction ensemble. This was done by using ProteinMPNN to design the first 83 residues of each chain in a monomeric context and using this “solubilized” version for design and prediction. Prior to ordering, solubilizing mutations were reverted to the original trimerization domain.

### Calculation of pae_interface

For a single AF2 prediction, contacts between chains are selected using the PyRosetta interface selector. Predicted aligned error (PAE) for each pair of residues in this selection is then extracted from the AF2 outputs and averaged to generate a pae_interface value for that prediction.

### Diffusion of symmetric icosahedra

RFDiffusion with symmetric conditioning was used to generate icosahedral backbones.^19^ Eight sequences for each backbone were designed with ProteinMPNN on the full 60-mer assembly using tied residues across each icosahedral protomer. Trimeric structure predictions for each designed sequence were generated with AF2 in single-sequence mode, using 12 recycles and an initial guess with a trimer derived from the design model. Final designs were filtered based on trimer backbone RMSD to the design-model trimer of less than 2 Å, AF2 pLDDT greater than 85, and AF2 pAE less than 10.

### Protein expression and purification

DNA for all designs was ordered with a verified sequence cloned into a pET29b+ plasmid from IDT. DNA was transformed into BL21 *E. coli* and directly transferred to autoinduction cultures after transformation. Cultures were grown overnight at 37 °C, spun down, and lysed by sonication. For single-component assemblies lysis was performed in TBS with PMSF and DNase (50 mM Tris, 500 mM NaCl, 20 mM imidazole, 1 mM PMSF, and 10 mM DNAse). Lysates were clarified by centrifugation at 10,000 x g for 20 minutes, then applied to nickel-NTA columns equilibrated with lysis buffer. After binding, columns were washed with 2 CV of wash buffer (50 mM Tris, 500 mM NaCl, 20 mM imidazole). Proteins were eluted from columns using 1 CV of elution buffer (50 mM Tris, 500 mM NaCl, 500 mM imidazole). For two-component assemblies, 3-[(3-cholamidopropyl) dimethylammonio]-1-propanesulfonate (CHAPS) was added to all buffers at 0.75% w/v.

### Multi-component protein assembly

Designs composed of two or more proteins (P3 arrays, de novo 2D arrays and high T nanocages) were expressed separately and purified via SEC in TBS with glycerol, arginine and CHAPS (30 mM Tris, 300 mM NaCl, 500 mM arginine, 5% glycerol, 0.75% CHAPS). After purification, proteins were concentration matched and mixed in equimolar ratios. The mixture was then dialyzed into TBS (25 mM Tris, 150 mM NaCl) overnight using a 3 kDa molecular-weight-cutoff (MWCO) dialysis cassette.

### Fluorescent labeling and super-resolution microscopy

P3 nanotubes were assembled as described above and exposed lysine residues were sub-stoichiometrically labeled with Alexa Fluor 594 NHS Ester (Thermo Fisher). Unassembled P3 proteins were separately labeled with Alexa Fluor 488 NHS Ester (Thermo Fisher). After nanotube assembly, 488-labeled components were added to the nanotubes and allowed to assemble for 4 hours before imaging. Assembled nanotubes were diluted to ensure homogeneous distribution on the glass surface of an Ibidi 18-well µ-slide (Ibidi, cat# 81817). Nanotubes were allowed to stick to the glass surface for 1 hour prior to imaging on a commercial OMX-SR system (GE Healthcare). Toptica diode lasers with excitation at 488 nm and 546 nm were used. Emission was collected on two separate PCO.edge sCMOS cameras using an Olympus 60× 1.42NA PlanApochromat oil immersion lens. 512 × 512 images (pixel size 6.5 μm) were captured with no binning. Acquisition was controlled with AcquireSR Acquisition control software. Structured Illumination Z-stacks were collected with a step size of 125 nm and a 1 µm total volume. Reconstructions were processed SoftWoRx 7.0.0 (GE Healthcare) using a weiner filter of 0.001. 2-color Image Z-stacks were collected with a step size of 500 nm and a 3 µm total volume before max intensity projection in SoftWoRx. Images from different color channels were registered in SoftWoRx using parameters generated from a gold grid registration slide (GE Healthcare).

### Negative stain electron microscopy

Specimens were prepared by applying 3 µL of sample to glow-discharged 300-mesh carbon-coated copper grids (Electron Microscopy Sciences, CF-300-Cu-50; 30 s at 15 mA), waiting 30 seconds, blotting, applying 3 µL of 2% (w/v) uranyl formate stain, and repeating the staining procedure three times. Negative-stain data were collected on a 120 kV Talos L120C transmission electron microscope (Thermo Fisher Scientific) equipped with a BM-Ceta camera using a 1 s exposure time. For datasets processed for 2D classification and reconstruction, images were recorded at 57,000× magnification, corresponding to a pixel size of 2.49 Å, with defocus values ranging from −1.3 µm to −2.3 µm sampled in six steps. Data collection was automated using E Pluribus Unum (EPU; Thermo Fisher Scientific). Image processing was performed in cryoSPARC (v4.7.0 and v4.7.1; Structura Biotechnology Inc.). Images were imported with constant CTF and particles were picked with either blob picking or template picking. Particle picks were then manually inspected, extracted, and subjected to 2D classification. Classes were then manually inspected to identify distinct species, which were reconstructed separately into ab-initio models and refined with symmetry as described above.

### Cryo-electron microscopy sample preparation

Prior to plunge-freezing in liquid ethane, samples were applied to glow-discharged lacey carbon grids and blotted using a Vitrobot at 4 °C and 100% humidity. Samples were prepared at 50 µM concentration in 50 mM Tris 150 mM NaCl pH 8 buffer unless otherwise stated. Nanocage samples were prepared by adding 3 µL of sample with a 7s wait time, 3s blot time, and blot force 0. SL109 nanotubes and SL112-IP 2D- and 3D-arrays were applied and blotted by hand three times inside the humidified Vitrobot chamber, then blotted for 1s with blot force 0. Icos-redes was prepared in 50 mM Tris 300 mM NaCl 200 mM L-Arginine HCl 5% (v/v) glycerol, pH 8.0 at 10 µM concentration. Thirty seconds after sample application, grids were blotted for 1.5 s with a blot force of 0 at 25 °C.

### Cryo electron microscopy data collection

Cryo-EM data were collected using SerialEM on Thermo Fisher Scientific transmission electron microscopes equipped with Gatan K3 direct electron detectors operated in counting mode. Single-particle datasets for SL109 nanocages and nanotubes, as well as tilt series for SL109 nanotubes and SL112-IP, were collected using a 200 kV Glacios microscope. We collected a total of 4,185 movies for SL109 nanocages and a total of 423 movies for SL109 nanotubes. Tilt series were collected at 2.064 Å/pixel in 3° increments from −60° to +60°, with four frames per tilt, a total dose of 115 e⁻/Å², and a target defocus range of −2 to −4 µm. Single-particle datasets were collected at a pixel size of 0.89 Å/pixel, with a total dose of 50 e⁻/Å² and a target defocus range of −1 to −2 µm. SL110-T and SL112-D datasets were collected at the Pacific Northwest Center for Cryo-EM using a Krios microscope (Thermo Fisher Scientific), yielding 3,185 and 4,619 movies, respectively. Data were collected at a pixel size of 0.45 Å/pixel, with a total dose of 37.5 e⁻/Å² and a target defocus range of −1 to −2 µm. SL112-IP was collected at the University of Washington using a Krios (Thermo Fisher) 300 kV with a total of 2,647 movies at a pixel size of 0.835 Å/pixel with a total dose of 60 e⁻/Å² and a target defocus of-1 to-2 microns. Icos-og was collected at the University of Washington using a Krios (Thermo Fisher) 300 kV with a total of 3,700 movies at a pixel size of 0.835 Å/pixel with a total dose of 47 e⁻/Å² and a target defocus of-1 to-2 microns.

### Contact Energy (dgCon) measurements and calculations

Contact energy was determined as previously described.^20^ Briefly, SL108-SL112 were purified via SEC and incubated with 0.19% CHAPS overnight. SEC was then repeated to determine the fraction of the sample in assembled nanocages and disassembled to trimers. Contact energies were calculated from the concentrations of free trimers and assembled nanocages using equilibrium models.^33,34^

### Cryo electron tomography reconstruction

Tilt series were reconstructed into motion-corrected volumes using AreTomo3.^35^ Reconstructions were performed using default settings, with a z-volume of 2,200 pixels and binning by a factor of 8. Tomograms were then inspected in IMOD. The viewing plane was adjusted to align with objects of interest using the “slicer tool,” and individual slices were exported.

### Single particle cryo-EM data processing

All image processing was performed in cryoSPARC and cryoSPARC Live (Structura Biotechnology Inc.) For all datasets, movies were imported and processed using Patch Motion Correction and Patch CTF estimation.

#### P3 nanotubes

Particles were identified using blob-based picking, extracted to a 1,000 pixel box size binned to 200 pixels, and subjected to several rounds of 2D classification. Classes of nanotube edges and hexagonal lattice views with no overlapping patterns were selected to obtain 450,300 particles to generate an ab-initio model. The model was then non-uniformly refined to obtain a 9.1 Å global resolution.

#### Icos-og

Particles were identified using blob-based picking, extracted to a 524 pixel box size and subjected to several rounds of 2D classification. Classes of tetrahedral nanoparticles were selected to obtain 23,059 particles to generate three ab-initio classes using HR-HAIR.^36^ The largest class with 13,478 particles was locally refined with C1 symmetry and subsequently non-uniformly refined with tetrahedral symmetry to obtain a 4.59 Å global resolution.

#### SL109 nanocage

Particles were identified using blob-based picking, extracted to a 600 pixel box size, and subjected to several rounds of 2D classification. Obtained classes of octahedral particles were then used for a second round of template picking. Particles were extracted to a 450 pixel box size and 2D classified to obtain a set of 334,861 particles identified to be octahedral species. An ab-initio model with octahedral symmetry was then generated and refined non-uniformly to obtain a map with 3.9 Å global resolution. This map was then imported in ChimeraX and a single trimer-trimer contact was isolated and input as a volume for local refinement. Symmetry-expanded particles from the octahedral map were refined into this volume with C1 symmetry to obtain a local reconstruction with 3.84 Å resolution.

#### SL109 nanotubes

Exposures were manually curated to obtain 164 movies with visible nanotubes. Particles were identified using blob-based picking, extracted to a 600 pixel box size, and subjected to several rounds of 2D classification to obtain 47,563 particles. An ab-initio model was constructed with no symmetry applied and imported to ChimeraX to isolate the region of the map containing 3 neighboring trimers. Particles were then homogeneously refined a second time into the reduced volume with no symmetry applied. Local refinement produced a final map at 8.1 Å resolution.

#### SL112-D mixed species

Particles were identified using templates from a negative stain dataset and extracted at a 1,800 pixel box size binned to 400 pixels. Particles were then subjected to several rounds of 2D classification to obtain 84,985 particles including all nanocage species. Ab-initio reconstruction was then performed to construct 5 initial models with no symmetry applied. These models were then manually inspected and refined with O, I, D3, or D5 symmetry. Symmetric models were then used to heterogeneously refine the full particle set into 4 separate classes. Each class was then homogeneously refined and non-uniformly refined. 13,863 particles were classified and refined into an icosahedral model with 6.2 Å global resolution. 35,022 particles were classified and refined into a D3 model with 6.2 Å global resolution. 10,470 particles were classified and refined into a D5 model with 6.7 Å global resolution. Particles in each set were then symmetry expanded and locally refined into models isolating trimer-trimer contacts with no symmetry applied. This yielded a set of final models with 3.9 Å local resolution for the icosahedral model, 6.5 Å local resolution for the D5 model, and 5.5 Å local resolution for the D3 model.

#### SL110-T

Particles were identified using blob-based picking, extracted to a 800 pixel box size binned to 200 pixels, and subjected to several rounds of 2D classification. Obtained classes of tetrahedral particles were then used for a second round of template picking. Particles were extracted and 2D classified to obtain a set of 111,369 particles identified to be tetrahedral species. An ab-initio model with C1 symmetry was then generated and refined homogeneously with tetrahedral symmetry to obtain a map with 3.3 Å global resolution. This map was then imported in ChimeraX and a single trimer-trimer contact was isolated and input as a volume for local refinement. Symmetry-expanded particles from the tetrahedral map were refined into this volume with C1 symmetry to obtain a local reconstruction with 2.4 Å resolution.

#### SL112-IP

Particles were identified using templates from a negative stain dataset and extracted at an 800 pixel box size binned to 200 pixels. Particles were then subjected to several rounds of 2D classification to obtain 33,718 particles identified to be icosahedral species and extracted at full resolution. An ab-initio model with icosahedral symmetry was then generated and refined homogeneously with icosahedral symmetry to obtain a map with 5.5 Å global resolution. This map was then imported in ChimeraX and a single trimer-trimer contact was isolated and input as a volume for local refinement. Symmetry-expanded particles from the icosahedral map were refined into this volume with C1 symmetry to obtain a local reconstruction with 3.8 Å resolution.

### In silico alanine scanning

Interface residues were selected with the PyRosetta interface selector. Single and double alanine mutations to SL110 and SL112 were then predicted with AF2 as described above. For each sequence, mean pae_interface across a 5-model ensemble was calculated and compared to the parent design. Mutations that increased pae_interface were selected and ordered for experimental screening.

### Multistate design of D3 assemblies

For multistate design of two edges of emergent D3 assemblies, backbone conformations were determined from a 6.8 Å cryo-EM model and combined into a single PDB file. Interface residues for both assembly interfaces were then selected with the PyRosetta interface selector. This list of residues was used to designate a tied residues dictionary to input to ProteinMPNN. Sequences were then designed by ProteinMPNN and predicted with AF2. Predictions were then aligned to both state A and state B design models and filtered for sequences which predicted both states within 3 Å accuracy. Designs were then manually inspected prior to ordering.

### Stepwise design of multistate icosahedral assemblies

Backbone conformations for P6_32 and I32 geometries were generated with RPXdock. To make conformations compatible for tiling into high-T icosahedra, we restricted rotation of the trimeric component between the two docking campaigns. Docks were then filtered for at least 10 contacting residues across the assembly interface. Sequences were designed by ProteinMPNN for residues at the assembly interface and folded with AF2. Designs for the P6_32 architecture were generated and experimentally verified prior to I32 design. The working sequence of P6_32-01 was threaded onto trimer and dimer design models prior to I32 docking. I32 docks were filtered as described above, with the additional requirement that less than 10% of interface residues overlap between the two models. Residues in the I32 interface were then designed with ProteinMPNN and folded with AF2. Sequences which predicted the I32 structure within 3 Å accuracy were then selected for targeted alanine mutations. For each prediction, pairwise pae values for interface residues were ranked, and residue pairs in the lower quartile of the range were selected for alanine mutation. Alanine mutants were then folded with AF2 and designs which predicted both P6_32 and I32 conformations within 5 Å accuracy and had a mean pae_interface value greater than 8 across a 5-prediction ensemble were selected for manual inspection.

## Notes

### Competing Interest Statement

The authors have declared no competing interest.

