## Supplemental Information for "Discovering and designing multistate protein assemblies"

**Amino Acid Sequences.** Two components of a single design are separated by a colon.

>P3_og
MEELFKEHKIVAVLRANSREEAIEIALAVFAGGVHLIEITFTVPDADEVIKRLEMLKRAGAIIGAGTVTSVEQCREAVE
SGAEFIVSPHLDEEISQFCKEEGVFYMPGVMTPTELVKAMKLGHTILKLFPGEVVGPQFVEAMKGPFPNVKFVPTGGVN
LDNVCEWFEAGVLAVGVGSALVEGKPSEVAEKARRFVKKIRGCLEHHHHHH:MEELFKEHKIVAVLRANSVEEAISKAL
AVFAGGVHLIEITFTVPDADQVIKELEFLKEAGAIIGAGTVTSVEQCREAVESGAEFIVSFHLDEEISQFCKEEGVFYM
PGVMTPTELVKAMKLGHTILKLVPGEVVGPQFVEAMKGPFPNVKFVPTGGVNLDNVCEWFEAGVLAVGVGSALVEGEPA
EVAELAIRFVEKIEGCLEHHHHHH

>P3_redes1
MEELFKEHKIVAVLRANSVEEAIAIALAVFAGGVHLIEITFTVPDADEVIKRLEPLKKRGAIIGAGTVTSVEQCREAVE
SGAEFIVSPHLDEEISQFCKEEGVFYMPGVMTPTELVKAMKLGHTILKLFPGEVVGPQFVEAMKGPFPNVKFVPTGGVN
LDNVCEWFEAGVLAVGVGSALVEGSPAEVAKKAQRFVEKIRGCLEHHHHHH:MEELFKEHKIVAVLRANSVEEAIAKAL
AVFAGGVHLIEITFTVPDADQVIKDLEFLKERGAIIGAGTVTSVEQCREAVESGAEFIVSPHLDEEISQFCKEEGVFYM
PGVMTPTELVKAMKLGHTILKLVPGEVVGPQFVEAMKGPFPNVKFVPTGGVNLDNVCEWFEAGVLAVGVGSALVEGSPA
EVAKLAIEFVEKIEGCLEHHHHHH

>P3_redes2
MEELFKEHKIVAVLRANSVEEAIRIALAVFAGGVHLIEITFTVPDADEVIKRLEPLKKRGAIIGAGTVTSVEQCREAVE
SGAEFIVSPHLDEEISQFCKEEGVFYMPGVMTPTELVKAMKLGHTILKLFPGEVVGPQFVEAMKGPFPNVKFVPTGGVN
LDNVCEWFEAGVLAVGVGSALVEGSPAEVARKAQRFVEKIRGCLEHHHHHH:MEELFKEHKIVAVLRANSVEEAIRKAL
AVFAGGVHLIEITFTVPDADQVIKDLEFLKKRGAIIGAGTVTSVEQCREAVESGAEFIVSPHLDEEISQFCKEEGVFYM
PGVMTPTELVKAMKLGHTILKLVPGEVVGPQFVEAMKGPFPNVKFVPTGGVNLDNVCEWFEAGVLAVGVGSALVEGSPA
EVATLAIEFVEKIEGCLEHHHHHH

>P3_redes3
MEELFKEHKIVAVLRANSVDEAEAIALAVFAGGVHLIEITFTVPDADEVIKRLEPLKKRGAIIGAGTVTSVEQCREAVE
SGAEFIVSPHLDEEISQFCKEEGVFYMPGVMTPTELVKAMKLGHTILKLFPGEVVGPQFVEAMKGPFPNVKFVPTGGVN
LDNVCEWFEAGVLAVGVGSALVEGSPAEVAKKAMRFVEKIRGCLEHHHHHH:MEELFKEHKIVAVLRANSVDEAIAKAL
AVFAGGVHLIEITFTVPDADQVIKDLEFLKKRGAIIGAGTVTSVEQCREAVESGAEFIVSPHLDEEISQFCKEEGVFYM
PGVMTPTELVKAMKLGHTILKLVPGEVVGPQFVEAMKGPFPNVKFVPTGGVNLDNVCEWFEAGVLAVGVGSALVEGSPS
EVAELAIKFVEKIEGCLEHHHHHH

>P3_redes4
MEELFKEHKIVAVLRANSVDEAVAIALAVFAGGVHLIEITFTVPDADEVIKRLEPLKERGAIIGAGTVTSVEQCREAVE
SGAEFIVSPHLDEEISQFCKEEGVFYMPGVMTPTELVKAMKLGHTILKLFPGEVVGPQFVEAMKGPFPNVKFVPTGGVN
LDNVCEWFEAGVLAVGVGSALVEGSPAEVAEKAMRFVEKIRGCLEHHHHHH:MEELFKEHKIVAVLRANSVDEAIAKAL
AVFAGGVHLIEITFTVPDADQVIKDLEFLKKRGAIIGAGTVTSVEQCREAVESGAEFIVSPHLDEEISQFCKEEGVFYM
PGVMTPTELVKAMKLGHTILKLVPGEVVGPQFVEAMKGPFPNVKFVPTGGVNLDNVCEWFEAGVLAVGVGSALVEGSPA
EVAELAIKFVKKIEGCLEHHHHHH

>icos_og
MEEEEELKEELIELNEKALKYPENAFFYTLLALGLIAEKLGLKGPDIYKGVYQFKSVEFDEEQKDAIKKVFLYLLQLAR
DPRVHFAATDIILQFLLFLALLEHHHHHH

>icos_redes
MEEEEELKEELIELNEKALKYPENAFFYTLLALGLIAEKLGIPVEKVVRGLFELMEGKATEEQKDAIKKVFLYLLQLAR
DPRVHFAATDIILQFLLFLALLEHHHHHH

>SL108
PELFLQDLRSLVEAARILARLARQRGDEHALERAARWAEQAARQAERLARQARKEGNLELALKALQILVNAAYVLAEIA
RDRGNEELLEYAARLAEEAARQAIEIWAQAMEEGNQQLRTKAAHIILRAAEVLLEIARDRGNQELLEKAASLVDAVAAL
QQAAAAILEGDVEKAVRAAQEAVKAAKEAGDNDMLRAVAIAALRIAKEAEKQGNVEVAVKAARVAVEAAKQAGDQDVLA
KVAIQAARIMTEALKQGNLEVALEAAKVASEALKQTLEHHHHHH
>SL109
PELFLQDLRSLVEAARILARLARQRGDEHALERAARWAEQAARQAERLARQARKEGNLELALKALQILVNAAYVLAEIA
RDRGNEELLEYAARLAEEAARQAIEIWAQAMEEGNQQLRTKAAHIILRAAEVLLEIARDRGNQELLEKAASLVDAVAAL
QQAAAAILEGDVEKAVRAAQEAVKAAKEAGDNDMLRAVAIAALRIAKEAEKQGNVEVAVKAARVAVEAAKQAGDQDVLA
KVADQAYRIAHEALKQGNLEVALEASKVAMEALKQTLEHHHHHH
>SL110
PELFLQDLRSLVEAARILARLARQRGDEHALERAARWAEQAARQAERLARQARKEGNLELALKALQILVNAAYVLAEIA
RDRGNEELLEYAARLAEEAARQAIEIWAQAMEEGNQQLRTKAAHIILRAAEVLLEIARDRGNQELLEKAASLVDAVAAL
QQAAAAILEGDVEKAVRAAQEAVKAAKEAGDNDMLRAVAIAALRIAKEAEKQGNVEVAVKAARVAVEAAKQAGDQDVLK
KVAAQAFRIALEALKQGNVEVALEALKVADEALKQTLEHHHHHH
>SL111
PELFLQDLRSLVEAARILARLARQRGDEHALERAARWAEQAARQAERLARQARKEGNLELALKALQILVNAAYVLAEIA
RDRGNEELLEYAARLAEEAARQAIEIWAQAMEEGNQQLRTKAAHIILRAAEVLLEIARDRGNQELLEKAASLVDAVAAL
QQAAAAILEGDVEKAVRAAQEAVKAAKEAGDNDMLRAVAIAALRIAKEAEKQGNVEVAVKAARVAVEAAKQAGDQDVLE
KVATQALRIAKEANKQGNVEVALEAIKVAQEALKQTLEHHHHHH
>SL112
PELFLQDLRSLVEAARILARLARQRGDEHALERAARWAEQAARQAERLARQARKEGNLELALKALQILVNAAYVLAEIA
RDRGNEELLEYAARLAEEAARQAIEIWAQAMEEGNQQLRTKAAHIILRAAEVLLEIARDRGNQELLEKAASLVDAVAAL
QQAAAAILEGDVEKAVRAAQEAVKAAKEAGDNDMLRAVAIAALRIAKEAEKQGNVEVAVKAARVAVEAAKQAGDQDVLA
KVAEQALRIAIEAIKQGNTEVAAEAIKVADEALKQTLEHHHHHH
>SL110-T
PELFLQDLRSLVEAARILARLARQRGDEHALERAARWAEQAARQAERLARQARKEGNLELALKALQILVNAAYVLAEIA
RDRGNEELLEYAARLAEEAARQAIEIWAQAMEEGNQQLRTKAAHIILRAAEVLLEIARDRGNQELLEKAASLVDAVAAL
QQAAAAILEGDVEKAVRAAQEAVKAAKEAGDNDMLRAVAIAALRIAKEAEKQGNVEVAVKAARVAVEAAKQAGDQDVLK
KVAAQAFRIALEAAKQGNVEVALEAAKVADEALKQTLEHHHHHH
>SL110-H
PELFLQDLRSLVEAARILARLARQRGDEHALERAARWAEQAARQAERLARQARKEGNLELALKALQILVNAAYVLAEIA
RDRGNEELLEYAARLAEEAARQAIEIWAQAMEEGNQQLRTKAAHIILRAAEVLLEIARDRGNQELLEKAASLVDAVAAL
QQAAAAILEGDVEKAVRAAQEAVKAAKEAGDNDMLRAVAIAALRIAKEAEKQGNVEVAVKAARVAVEAAKQAGDQDVLK
KVAAQAFRIAAEALKQGNVEVALEALKVADEAAKQTLEHHHHHH
>Sl112-D
PELFLQDLRSLVEAARILARLARQRGDEHALERAARWAEQAARQAERLARQARKEGNLELALKALQILVNAAYVLAEIA
RDRGNEELLEYAARLAEEAARQAIEIWAQAMEEGNQQLRTKAAHIILRAAEVLLEIARDRGNQELLEKAASLVDAVAAL
QQAAAAILEGDVEKAVRAAQEAVKAAKEAGDNDMLRAVAIAALRIAKEAEKQGNVEVAVKAARVAVEAAKQAGDQDVLA
KVAEQALRIAIEAIKQGNTEVAAEAIKVADEAAKQTLEHHHHHH
>SL112-IP
PELFLQDLRSLVEAARILARLARQRGDEHALERAARWAEQAARQAERLARQARKEGNLELALKALQILVNAAYVLAEIA
RDRGNEELLEYAARLAEEAARQAIEIWAQAMEEGNQQLRTKAAHIILRAAEVLLEIARDRGNQELLEKAASLVDAVAAL
QQAAAAILEGDVEKAVRAAQEAVKAAKEAGDNDMLRAVAIAALRIAKEAEKQGNVEVAVKAARVAVEAAKQAGDQDVLA
KVAEQALRIAAEAIKQGNTEVAAEAIKVADEAAKQTLEHHHHHH

>D3-1
PELFLQDLRSLVEAARILARLARQRGDEHALERAARWAEQAARQAERLARQARKEGNLELALKALQILVNAAYVLAEIA
RDRGNEELLEYAARLAEEAARQAIEIWAQAMEEGNQQLRTKAAHIILRAAEVLLEIARDRGNQELLEKAASLVDAVAAL
QQAAAAILEGDVEKAVRAAQEAVKAAKEAGDNDMLRAVAIAALRIAKEAEKQGNVEVAVKAARVAVEAAKQAGDQDVLA
KVAEQAERIAKEALEQGNLEVAIEAIKVALEAKEQTLEHHHHHH
>D3-2
PELFLQDLRSLVEAARILARLARQRGDEHALERAARWAEQAARQAERLARQARKEGNLELALKALQILVNAAYVLAEIA
RDRGNEELLEYAARLAEEAARQAIEIWAQAMEEGNQQLRTKAAHIILRAAEVLLEIARDRGNQELLEKAASLVDAVAAL
QQAAAAILEGDVEKAVRAAQEAVKAAKEAGDNDMLRAVAIAALRIAKEAEKQGNVEVAVKAARVAVEAAKQAGDQDVLA
KVAEQAERIAKEALEQGNVEVAIEALKVALEAKEQTLEHHHHHH
>D3-3
PELFLQDLRSLVEAARILARLARQRGDEHALERAARWAEQAARQAERLARQARKEGNLELALKALQILVNAAYVLAEIA
RDRGNEELLEYAARLAEEAARQAIEIWAQAMEEGNQQLRTKAAHIILRAAEVLLEIARDRGNQELLEKAASLVDAVAAL
QQAAAAILEGDVEKAVRAAQEAVKAAKEAGDNDMLRAVAIAALRIAKEAEKQGNVEVAVKAARVAVEAAKQAGDQDVLA
KVAEQALRIAKEAIEQGNVEVGIEALKVALEAKKQTLEHHHHHH
>D3-4
PELFLQDLRSLVEAARILARLARQRGDEHALERAARWAEQAARQAERLARQARKEGNLELALKALQILVNAAYVLAEIA
RDRGNEELLEYAARLAEEAARQAIEIWAQAMEEGNQQLRTKAAHIILRAAEVLLEIARDRGNQELLEKAASLVDAVAAL
QQAAAAILEGDVEKAVRAAQEAVKAAKEAGDNDMLRAVAIAALRIAKEAEKQGNVEVAVKAARVAVEAAKQAGDQDVLA
KVAEQAKRIAIEAIKQGNVEVAIEALKVALEAEKQTLEHHHHHH
>D3-5
PELFLQDLRSLVEAARILARLARQRGDEHALERAARWAEQAARQAERLARQARKEGNLELALKALQILVNAAYVLAEIA
RDRGNEELLEYAARLAEEAARQAIEIWAQAMEEGNQQLRTKAAHIILRAAEVLLEIARDRGNQELLEKAASLVDAVAAL
QQAAAAILEGDVEKAVRAAQEAVKAAKEAGDNDMLRAVAIAALRIAKEAEKQGNVEVAVKAARVAVEAAKQAGDQDVLA
KVAEQARRIAIEAIKQGNVEVAIEALKVALEAEKQTLEHHHHHH
>D3-6
PELFLQDLRSLVEAARILARLARQRGDEHALERAARWAEQAARQAERLARQARKEGNLELALKALQILVNAAYVLAEIA
RDRGNEELLEYAARLAEEAARQAIEIWAQAMEEGNQQLRTKAAHIILRAAEVLLEIARDRGNQELLEKAASLVDAVAAL
QQAAAAILEGDVEKAVRAAQEAVKAAKEAGDNDMLRAVAIAALRIAKEAEKQGNVEVAVKAARVAVEAAKQAGDQDVLA
KVAEQAKRIAKEALEQGNVKVAIEAAKVAEEAEKQTLEHHHHHH
>D3-7
PELFLQDLRSLVEAARILARLARQRGDEHALERAARWAEQAARQAERLARQARKEGNLELALKALQILVNAAYVLAEIA
RDRGNEELLEYAARLAEEAARQAIEIWAQAMEEGNQQLRTKAAHIILRAAEVLLEIARDRGNQELLEKAASLVDAVAAL
QQAAAAILEGDVEKAVRAAQEAVKAAKEAGDNDMLRAVAIAALRIAKEAEKQGNVEVAVKAARVAVEAAKQAGDQDVLA
KVAEQALRIAEEALKQGNVEVAIEAAKVALEAKEQTLEHHHHHH
>D3-8
PELFLQDLRSLVEAARILARLARQRGDEHALERAARWAEQAARQAERLARQARKEGNLELALKALQILVNAAYVLAEIA
RDRGNEELLEYAARLAEEAARQAIEIWAQAMEEGNQQLRTKAAHIILRAAEVLLEIARDRGNQELLEKAASLVDAVAAL
QQAAAAILEGDVEKAVRAAQEAVKAAKEAGDNDMLRAVAIAALRIAKEAEKQGNVEVAVKAARVAVEAAKQAGDQDVLA
KVAEQAERIAIEAIKQGNVEVGIEAAKVALEAKKQTLEHHHHHH
>D3-9
PELFLQDLRSLVEAARILARLARQRGDEHALERAARWAEQAARQAERLARQARKEGNLELALKALQILVNAAYVLAEIA
RDRGNEELLEYAARLAEEAARQAIEIWAQAMEEGNQQLRTKAAHIILRAAEVLLEIARDRGNQELLEKAASLVDAVAAL
QQAAAAILEGDVEKAVRAAQEAVKAAKEAGDNDMLRAVAIAALRIAKEAEKQGNVEVAVKAARVAVEAAKQAGDQDVLA
KVAEQAERIAIEAIKQGNVEVGIEAAKVALEAKEQTLEHHHHHH

>D3-10
PELFLQDLRSLVEAARILARLARQRGDEHALERAARWAEQAARQAERLARQARKEGNLELALKALQILVNAAYVLAEIA
RDRGNEELLEYAARLAEEAARQAIEIWAQAMEEGNQQLRTKAAHIILRAAEVLLEIARDRGNQELLEKAASLVDAVAAL
QQAAAAILEGDVEKAVRAAQEAVKAAKEAGDNDMLRAVAIAALRIAKEAEKQGNVEVAVKAARVAVEAAKQAGDQDVLA
KVAEQAKRIAIEAIKQGNVEVGLEALKVALEAEKQTLEHHHHHH

>D3-11
PELFLQDLRSLVEAARILARLARQRGDEHALERAARWAEQAARQAERLARQARKEGNLELALKALQILVNAAYVLAEIA
RDRGNEELLEYAARLAEEAARQAIEIWAQAMEEGNQQLRTKAAHIILRAAEVLLEIARDRGNQELLEKAASLVDAVAAL
QQAAAAILEGDVEKAVRAAQEAVKAAKEAGDNDMLRAVAIAALRIAKEAEKQGNVEVAVKAARVAVEAAKQAGDQDVLA
KVAEQALRIAKEALEQGNVEVAIEAIKVALEAKKQTLEHHHHHH

>D3-12
PELFLQDLRSLVEAARILARLARQRGDEHALERAARWAEQAARQAERLARQARKEGNLELALKALQILVNAAYVLAEIA
RDRGNEELLEYAARLAEEAARQAIEIWAQAMEEGNQQLRTKAAHIILRAAEVLLEIARDRGNQELLEKAASLVDAVAAL
QQAAAAILEGDVEKAVRAAQEAVKAAKEAGDNDMLRAVAIAALRIAKEAEKQGNVEVAVKAARVAVEAAKQAGDQDVLA
KVAEQALRIAKEALEQGNVEVAIEAIKVAKEAKEQTLEHHHHHH

>D3-13
PELFLQDLRSLVEAARILARLARQRGDEHALERAARWAEQAARQAERLARQARKEGNLELALKALQILVNAAYVLAEIA
RDRGNEELLEYAARLAEEAARQAIEIWAQAMEEGNQQLRTKAAHIILRAAEVLLEIARDRGNQELLEKAASLVDAVAAL
QQAAAAILEGDVEKAVRAAQEAVKAAKEAGDNDMLRAVAIAALRIAKEAEKQGNVEVAVKAARVAVEAAKQAGDQDVLA
KVAEQAERIAIEAIKQGNVEVGIEALKVALEAKKQTLEHHHHHH

>D3-14
PELFLQDLRSLVEAARILARLARQRGDEHALERAARWAEQAARQAERLARQARKEGNLELALKALQILVNAAYVLAEIA
RDRGNEELLEYAARLAEEAARQAIEIWAQAMEEGNQQLRTKAAHIILRAAEVLLEIARDRGNQELLEKAASLVDAVAAL
QQAAAAILEGDVEKAVRAAQEAVKAAKEAGDNDMLRAVAIAALRIAKEAEKQGNVEVAVKAARVAVEAAKQAGDQDVLA
KVAEQAERIAIEAIKQGNVEVGIEALKVALEAKEQTLEHHHHHH

>D3-15
PELFLQDLRSLVEAARILARLARQRGDEHALERAARWAEQAARQAERLARQARKEGNLELALKALQILVNAAYVLAEIA
RDRGNEELLEYAARLAEEAARQAIEIWAQAMEEGNQQLRTKAAHIILRAAEVLLEIARDRGNQELLEKAASLVDAVAAL
QQAAAAILEGDVEKAVRAAQEAVKAAKEAGDNDMLRAVAIAALRIAKEAEKQGNVEVAVKAARVAVEAAKQAGDQDVLA
KVAEQAKRIAKEAIEQGNVEVGIEALKVALEAEKQTLEHHHHHH

>p6-01
GAADARLAAAARAELERLKQLPGKPEKPTLPAGWTLDRQLGERLFTLSRTDGDATVIVRLTGEINPSETKILTLEVLIV
RDGKALVFDLEVVNGRFVIRAVRFEADGALALDQSANAQYQLSRRYQGPDLSTLPEEFLRALRYYLALRGINRRLAAFI
EAFSAYWDWENELAWLAEVAAFASLEHHHHHH:SVAEEIEKRAKKISKELKKEGKNPEWIEELQRAADKLEEVARRATS
SDAKEIAKRAVKIAEELAKQGSNPKWIAELLLAAAKLVEVAARATSSVARWIAKVAVEAAELLAWAGENPKEIAELLLE
AAALVERAARATSPEEKNAALAEAIELLQEAARKAVEAAEERLERVKEASKRGVSSDQLKDLIRELAEIIETLIKIIRL
SNEAIKELIKALSLEHHHHHH

>multi-icos_parent
GAADARLAAAARAELERLKQLPGKPEKPTLPAGWRLHRQLGERLFTLSRRDGEATVIVRLTGEINPSETKILTLEVLIV
RDGKALVFDLEVVNGRFVIRAVRFEADGWLALDQSANAQYQLSRRYQGPDLSTLPEEFLRALRYYLALRGINRRLAAFI
EAFSAYWDWENELAWLAEVAAFASLEHHHHHH:SVAEEIEKRARKIAEELLAEGKNIEWVKELLRAADKLEEVARRATS
SDAKEIAKRAVKIAEELAKQGSNPKWIAELLLAAAKLVEVAARATSSVARWIAKVAVEAAELLAWAGENPKEIAELLLE
AAALVERAARATSPEEKNAALAEAIELLQEAARKAVEAAEERLERVKEASKRGVSSDQLKDLIRELAEIIETLIKIIRL
SNEAIKELIKALSLEHHHHHH

>multi-icos 110

```
GAADARLAAAARAELERLKQLPGKPEKPTLPAGWRLHRQLGERLFTLARRDGEATVIVRLTGEINPSETKILTLEVLIV
RDGKALVFDLEVVNGRFVIRAVRFEADGWLALDQSANAQYQLSRRYQGPDLSTLPEEFLRALRYYLALRGINRRLAAFI
EAFSAYWDWENELAWLAEVAAFASLEHHHHHH:SVAEEIEKRARKIAEELLAEGKNAEWVKELLRAADKLEEVARRATS
SDAKEIAKRAVKIAEELAKQGSNPKWIAELLLAAAKLVEVAARATSSVARWIAKVAVEAAELLAWAGENPKEIAELLLE
AAALVERAARATSPEEKNAALAEAIELLQEAARKAVEAAEERLERVKEASKRGVSSDQLKDLIRELAEIIETLIKIIRL
SNEAIKELIKALSLEHHHHHH
```

### Supplemental Figures

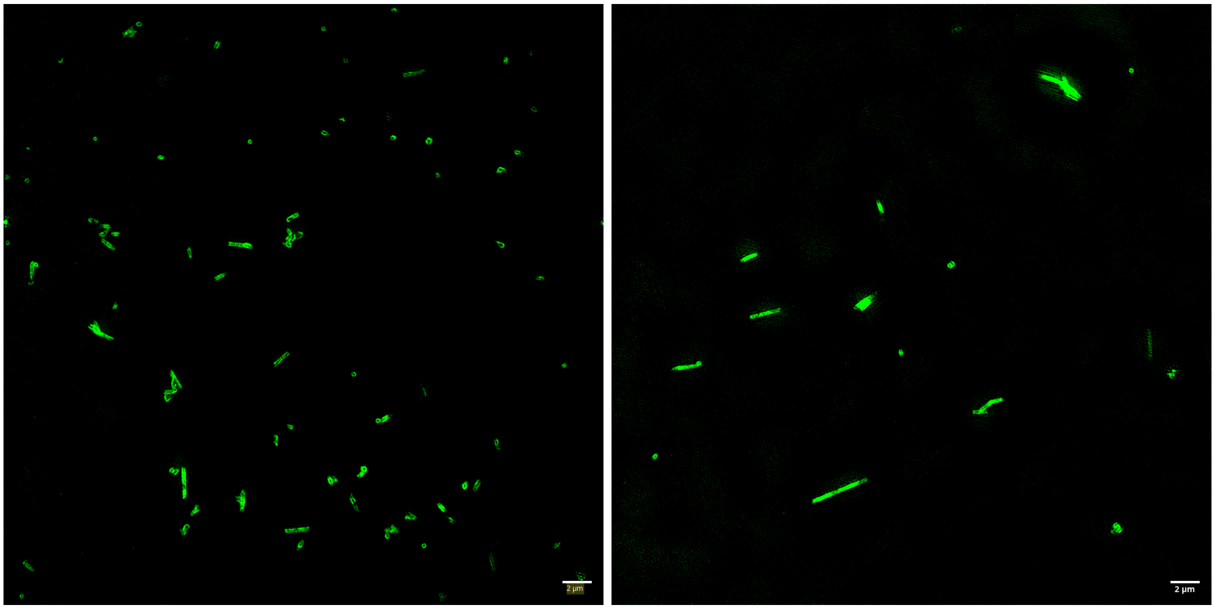


Figure S1: Super resolution of fluorescently labeled nanotubes. P3-og nanotubes were assembled and subsequently labeled with Alexa Fluor 488.

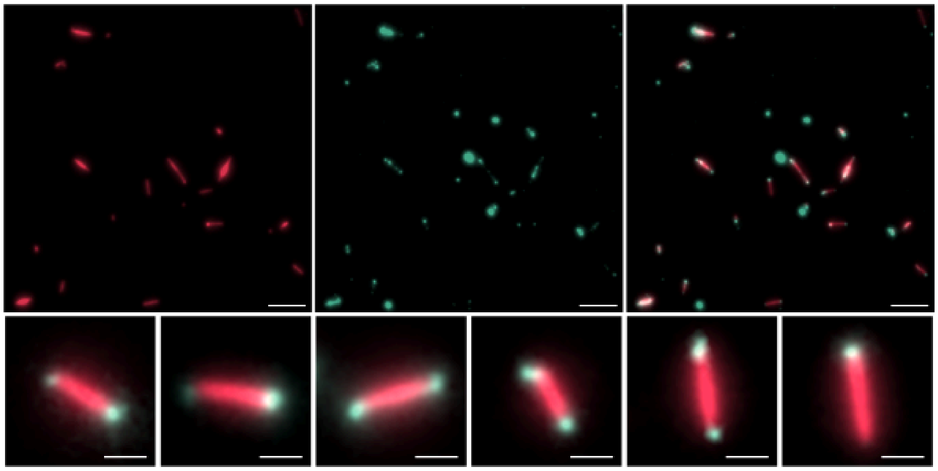

Figure S2: Nanotube seeds template the extension of additional proteins without slow assembly. Scale bar for full frame images is 10 microns, cutouts are 1 micron.

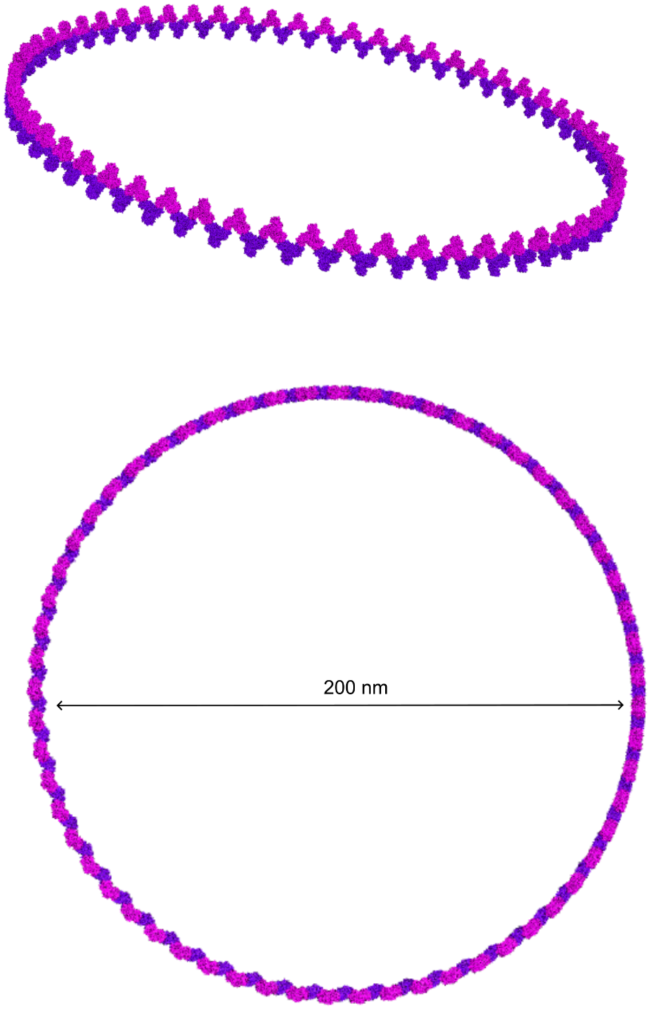


Figure S3: Model of a 200 nm ring formed by rotating P3-og trimers 3 degrees across the assembly interface.

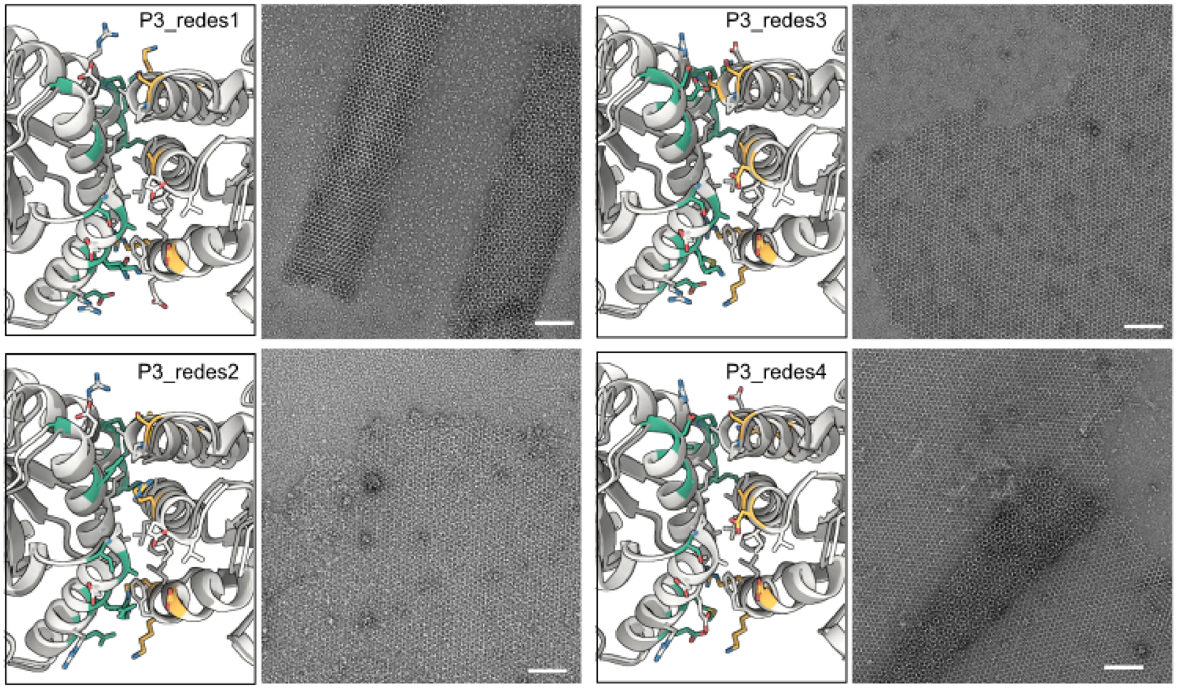


Figure S4: Interface designs and micrographs of P3 redesigns. Scale bar is 100 nm.

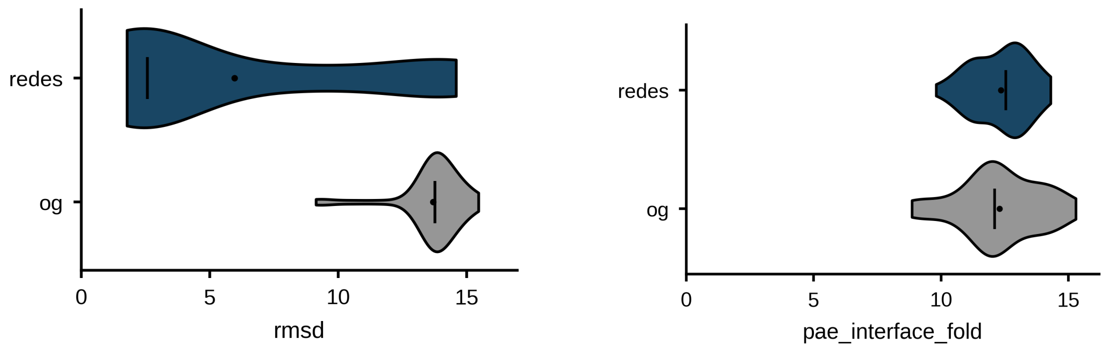


Figure S5: AF2 predictions of original (og) and redesigned (redes) diffused nanoparticles. Each sequence was predicted by AF2 multimer v3 25 times and pae_interface and RMSD to the icosahedral 2-fold were calculated for each prediction.

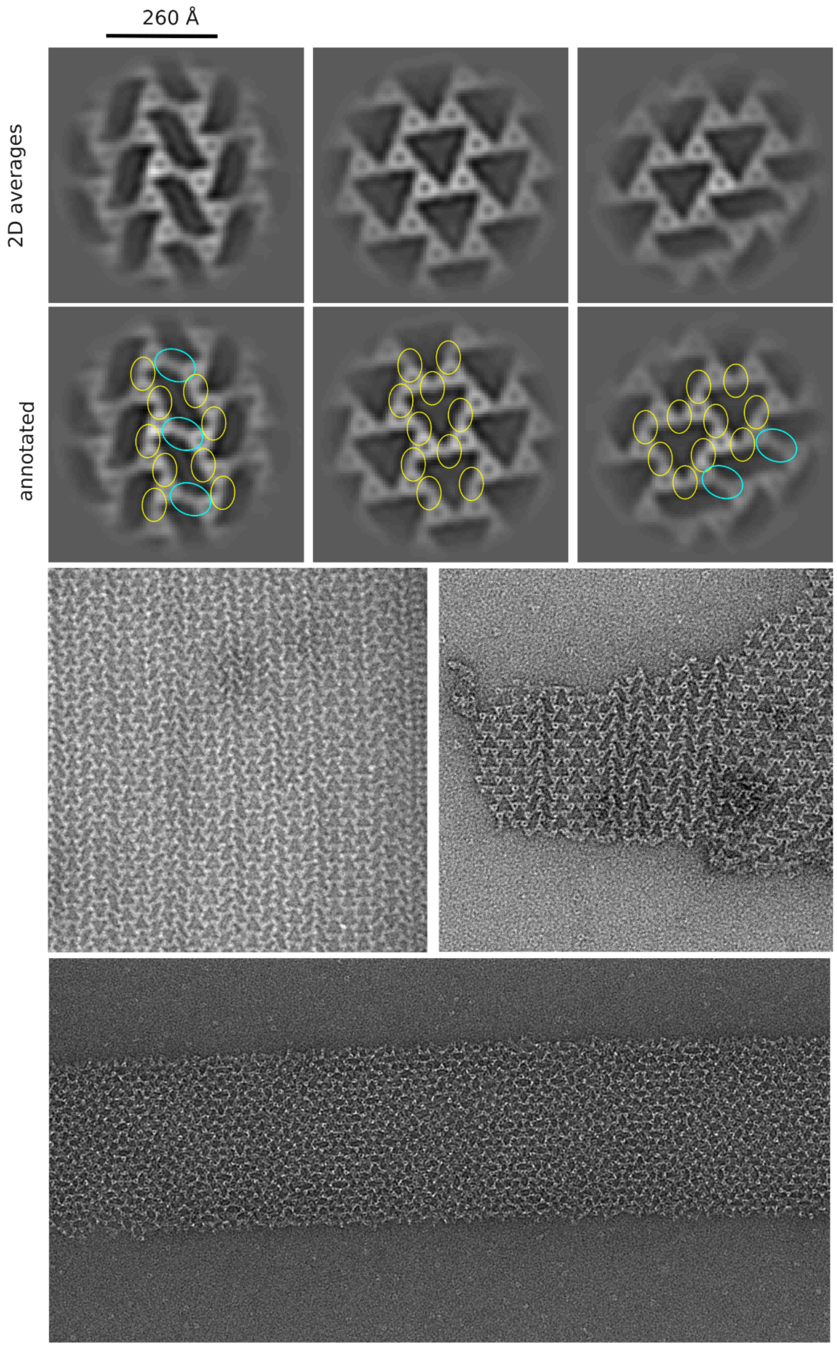


Figure S6: SL109 lattice geometry. 2D averages are annotated to show the arrangement of trimer-trimer contact modes within the lattice structure. S-curves show alternating yellow and blue contact modes, whereas triangular lattices only show yellow contact modes. Within large 2D arrays, triangles and S-curves exist in straight lines. Nanotubes show regularly repeating S-curve patterns.

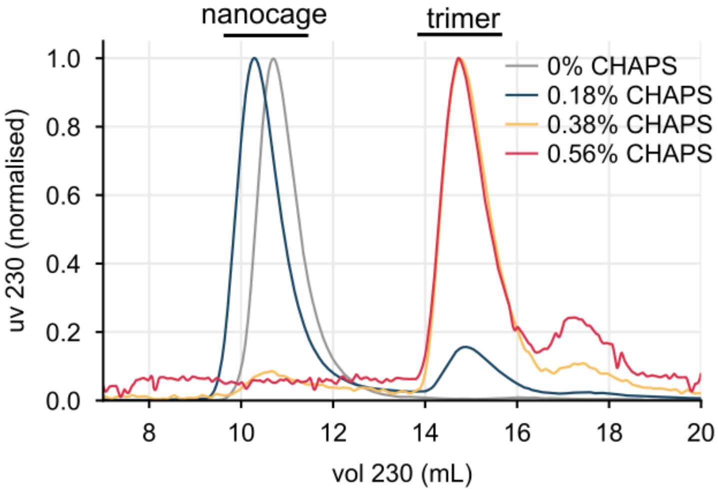


Figure S7: Representative SEC trace of nanocage disassembly with CHAPS treatment. Nanocages and trimers elute at 11 mL and 15 mL RT, respectively from a Superdex 200 column. Increasing concentrations of CHAPS shift the relative quantity of protein in the nanocage and trimer fractions.

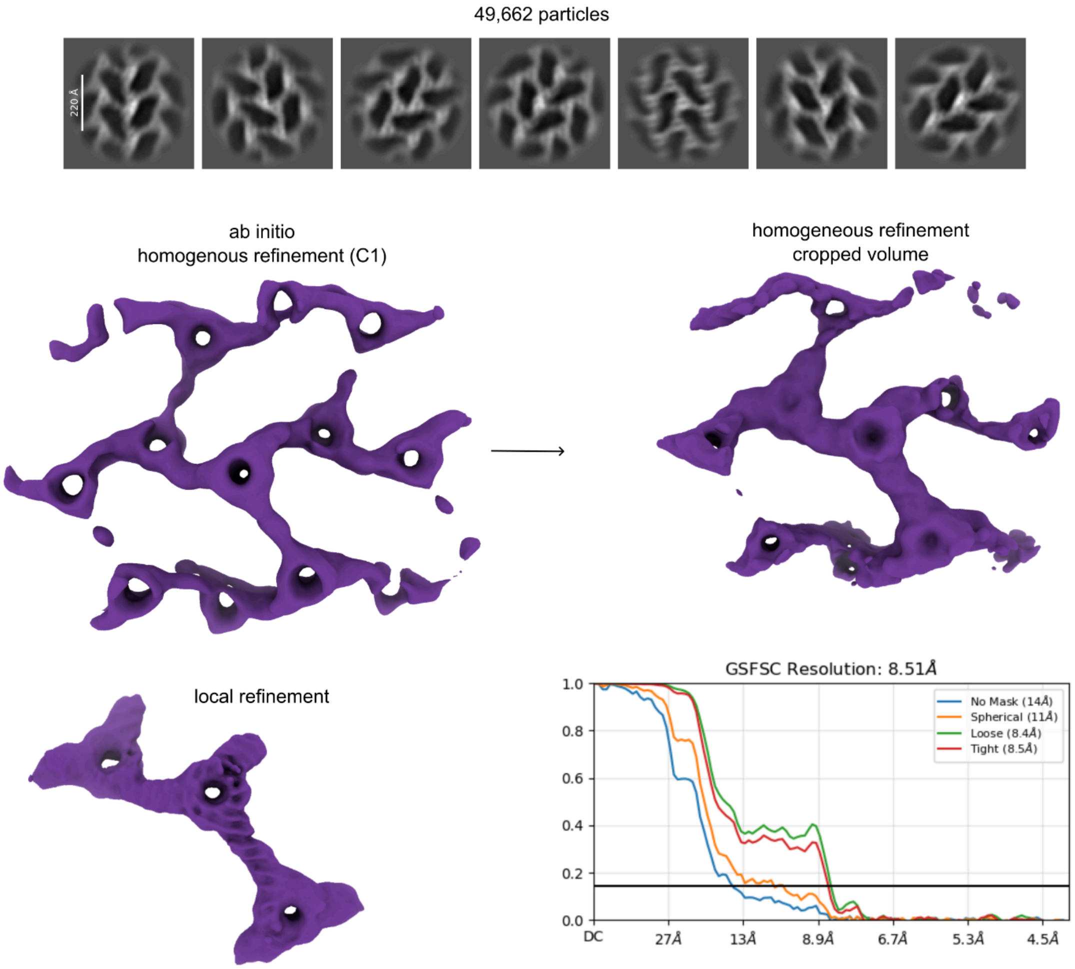


Figure S8: CryoEM processing of SL109 nanotubes. Particles in S-curved lattices were identified via 2D classification. These particles were used to generate an ab initio model capturing the full lattice geometry. This volume was then cropped in ChimeraX to focus on refining the particles into 3 neighboring trimers. Finally, a local refinement yielded a reconstruction at 8.5 Å resolution.

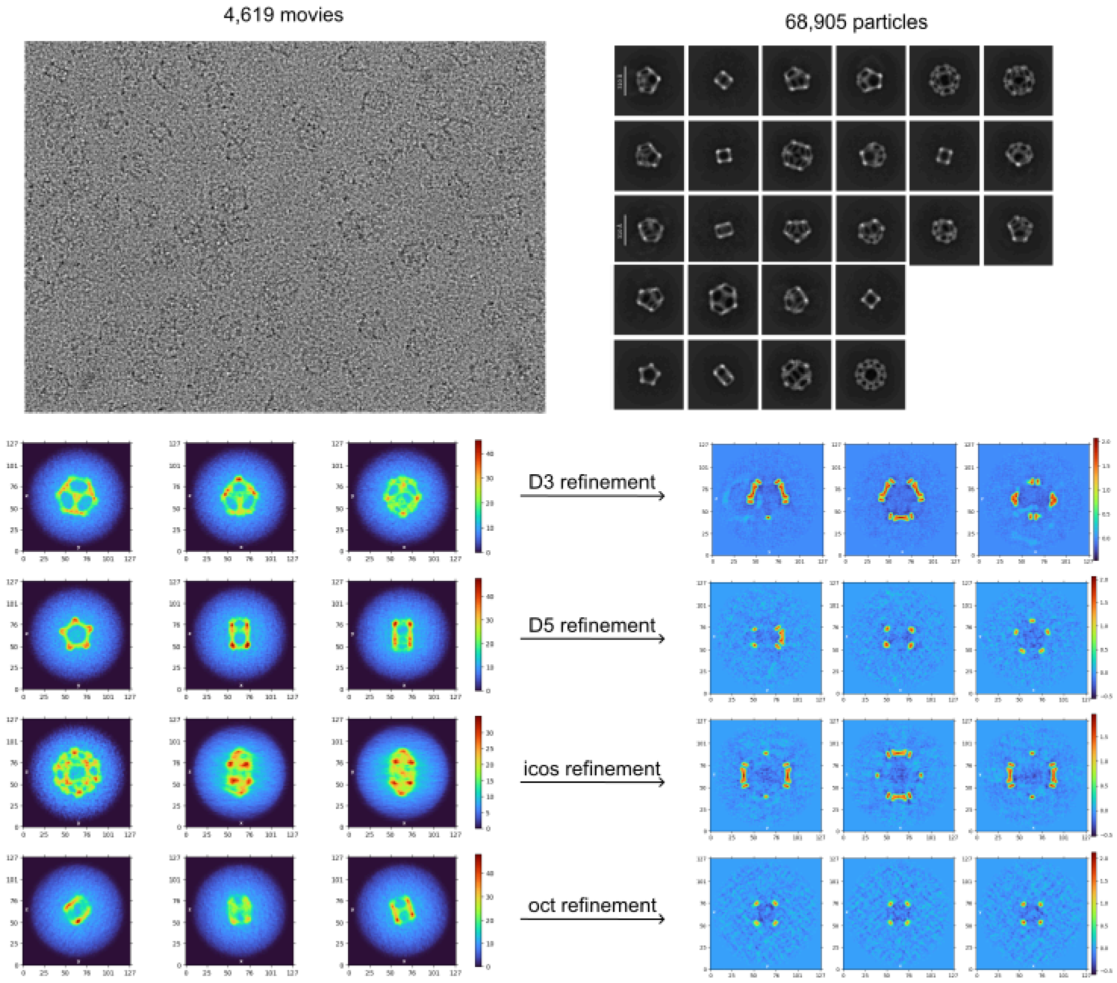


Figure S9: CryoEM processing of multiple species from SL112-D. Nanocage species were identified via 2D classification and 5 ab initio models were generated from the full nanocage series (1 junk class not shown). These ab initio volumes were manually inspected to identify the symmetry of each species and homogeneously refined to generate symmetric models for heterogeneous refinement. Downstream processing is described in detail in the Methods section.

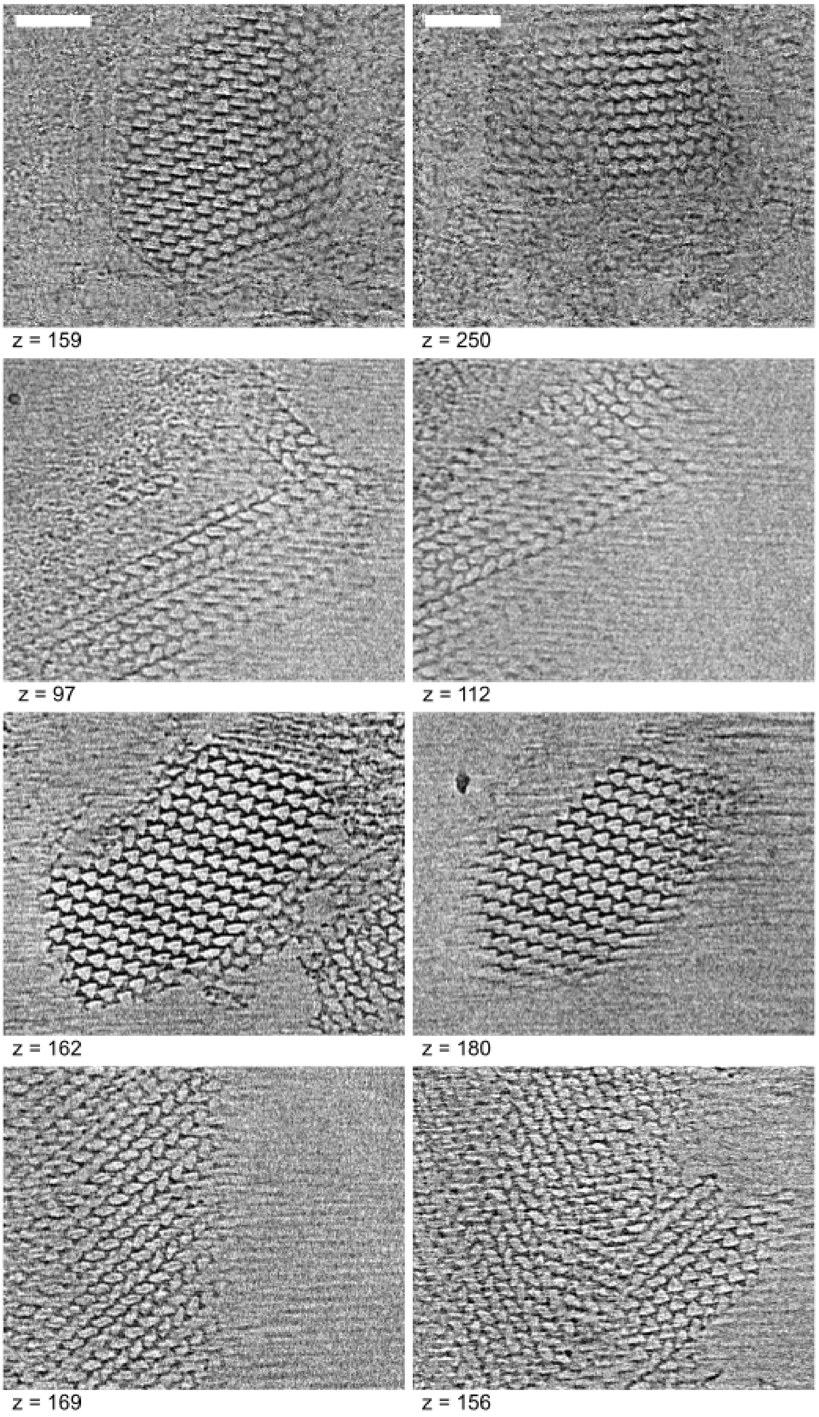


Figure S10: CryoET of SL112-IP 3D objects.
Tilt series were collected and reconstructed with AreTomo3. Slices from individual tomograms are arranged in rows.

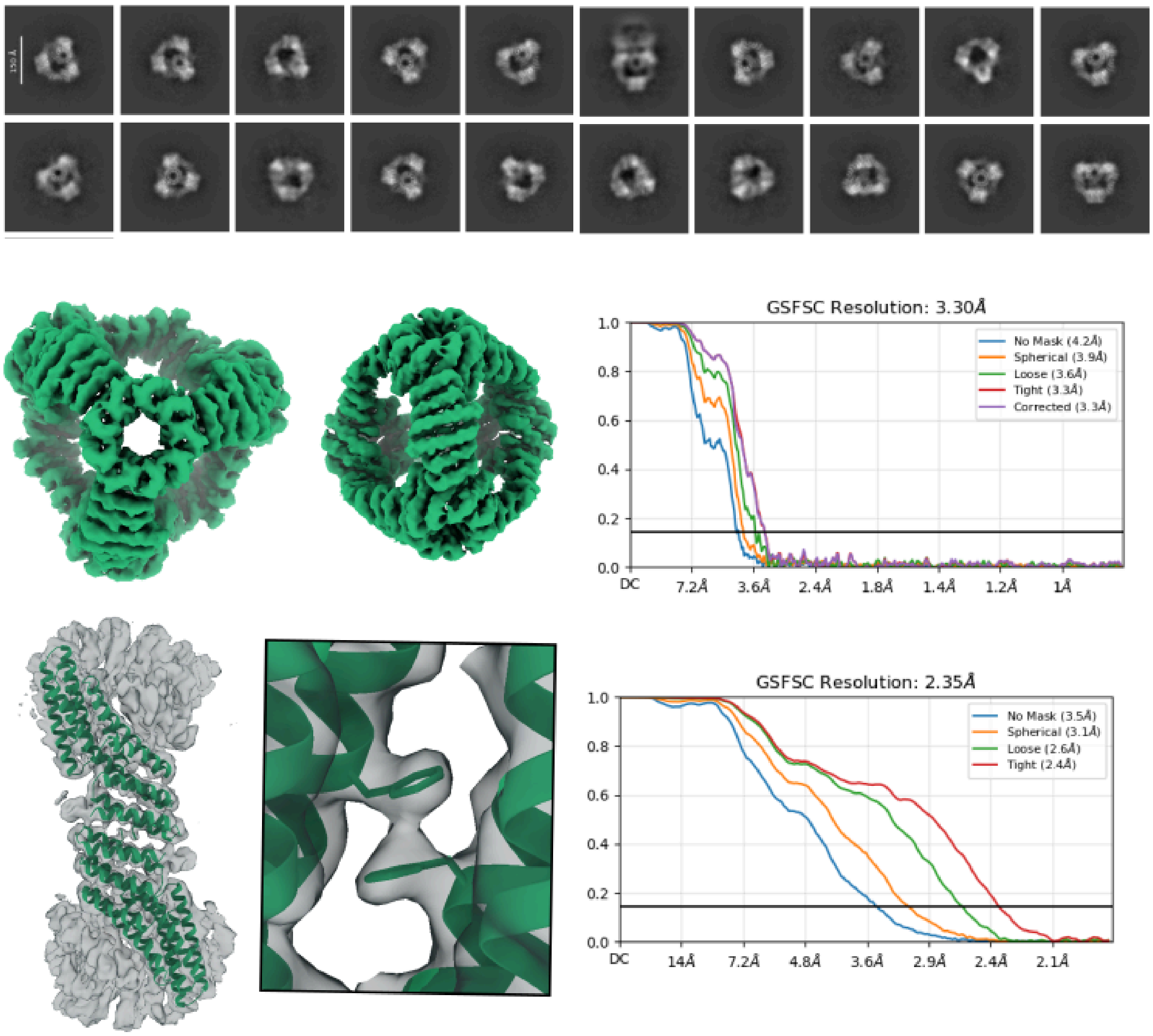


Figure S11: CryoEM processing of SL110-T tetrahedra. 2D classification showed many views of tetrahedral nanocages which were used to generate an ab initio model which was refined to 3.3 Å global resolution with tetrahedral symmetry applied. Particles were then symmetry expanded and refined into a local model of two contacting trimer chains at 2.4 Å resolution with C1 symmetry applied.

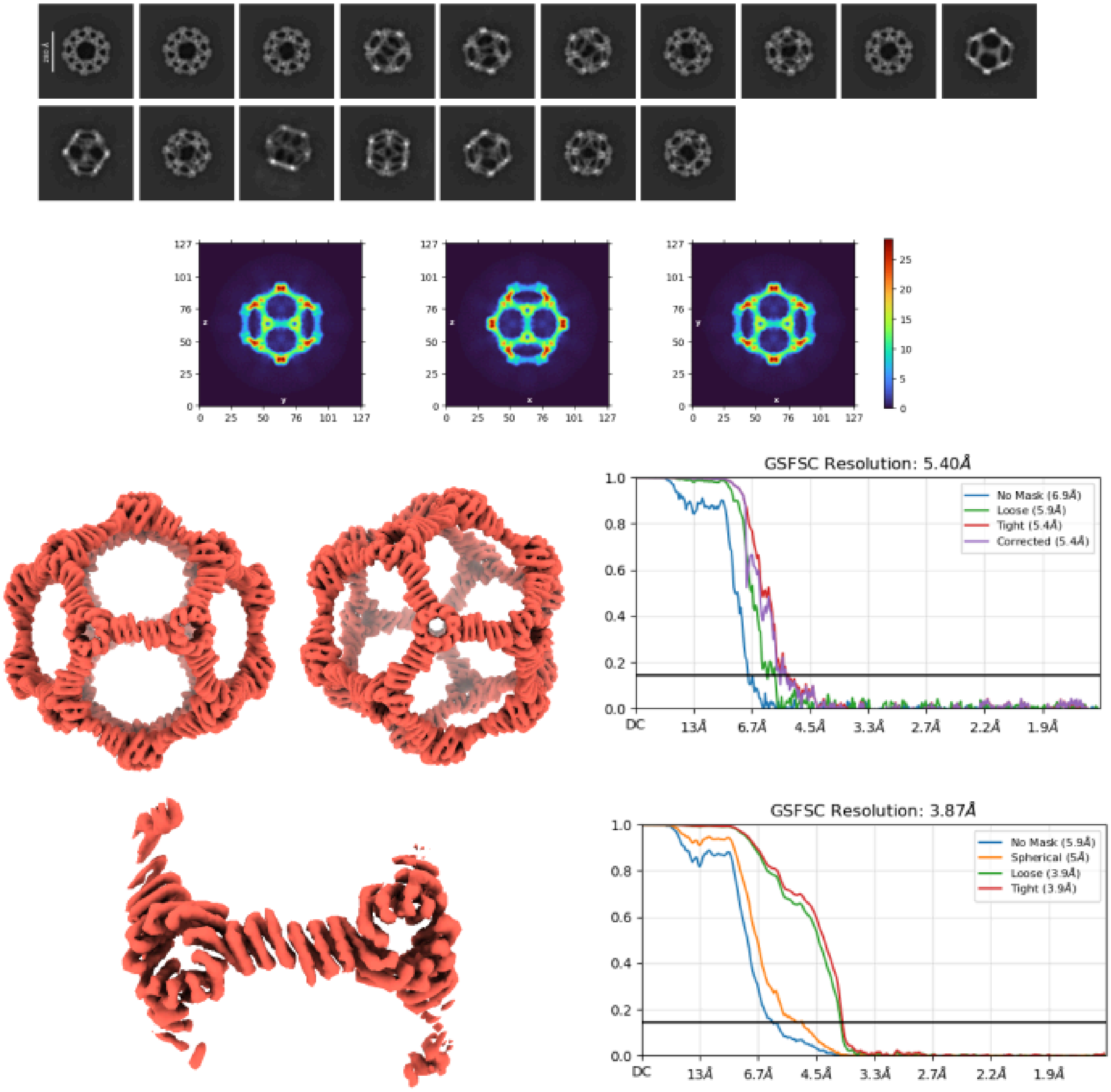


Figure S12: CryoEM processing of SL112-IP icosahedra. 2D classification showed many views of icosahedral nanocages which were used to generate an ab initio model which was refined to 5.4 Å global resolution with icosahedral symmetry applied. Particles were then symmetry expanded and refined into a local model of two contacting trimer chains at 3.9 Å resolution with C1 symmetry applied.

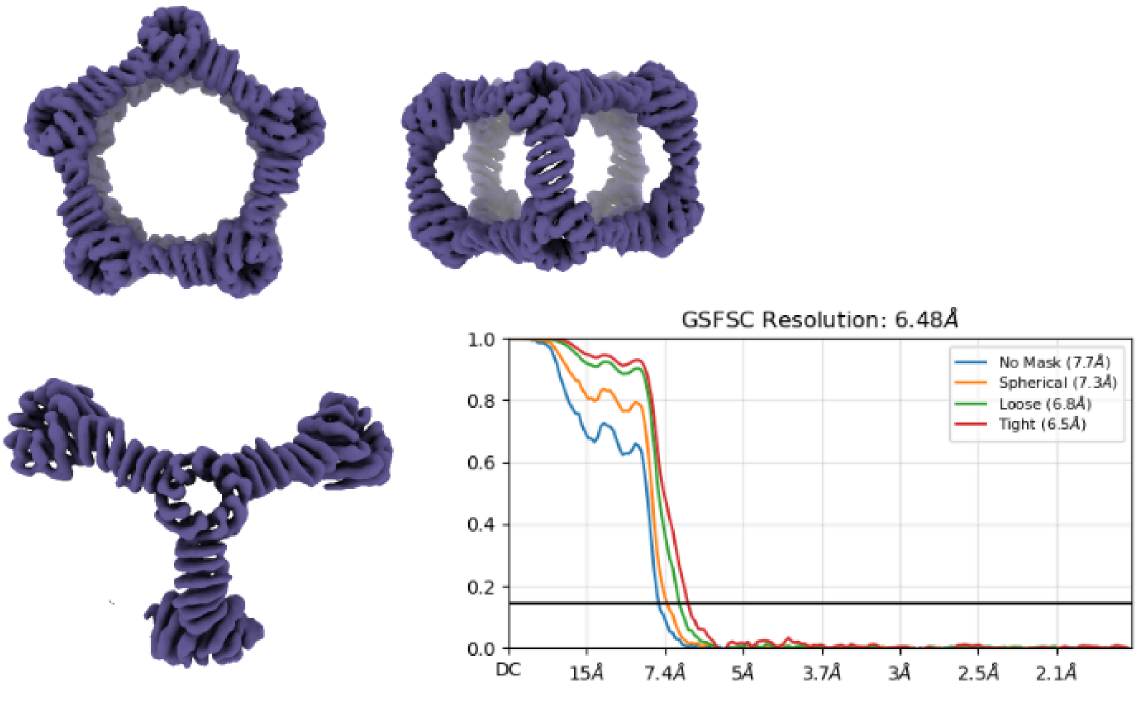


Figure S13: CryoEM processing of SL112-D D5 species. Particles were heterogeneously classified into several symmetric species as shown in Figure S8 and subsequently refined. Particles were then symmetry expanded and refined into a local model with both penton and 2-fold edges of the D5 cage at 6.5 Å resolution with C1 symmetry applied.

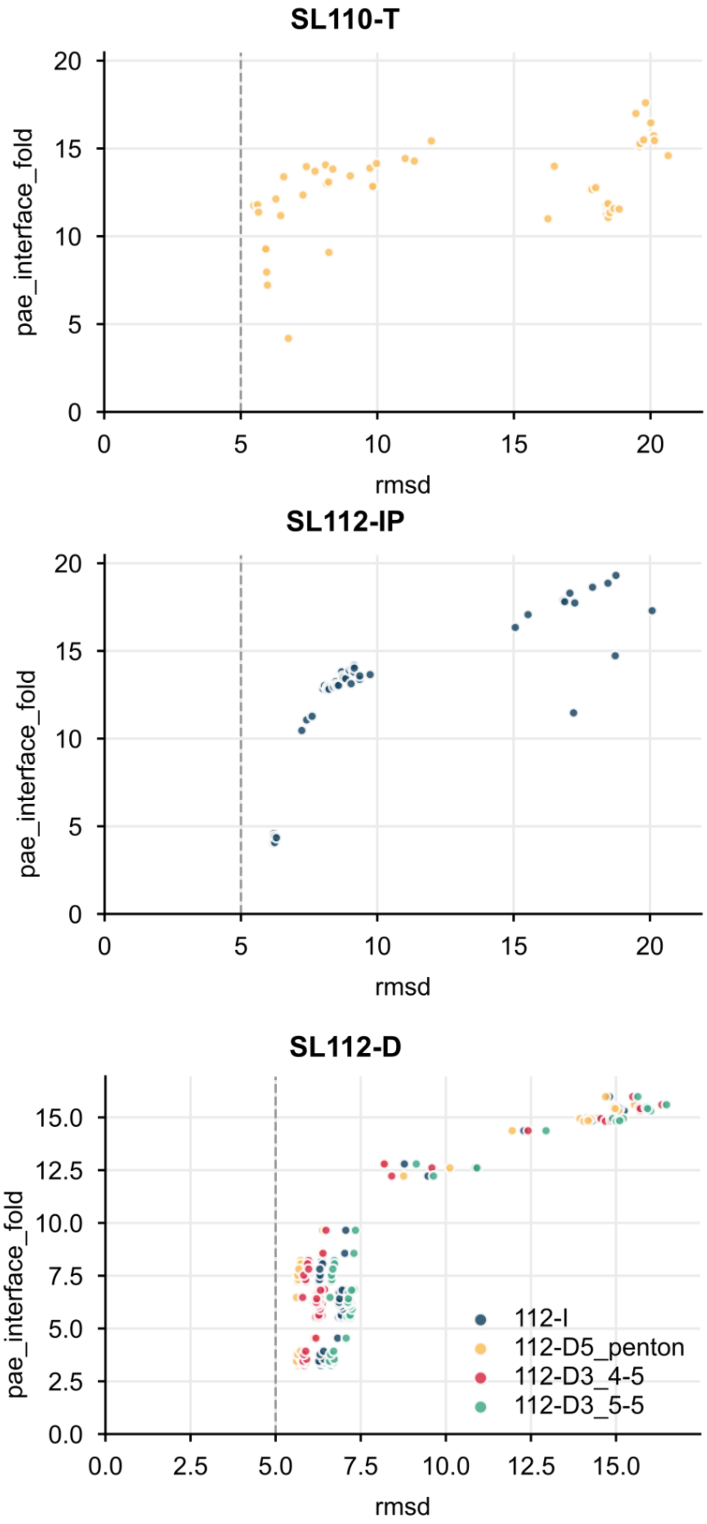


Figure S14: AF2 prediction of emergent assemblies. Sequences of SL110-T, SL112-IP and SL112-D were predicted by AF2 100 times and aligned to experimental cryo-EM models. RMSD to the experimental model and pae_interface were calculated for each prediction.

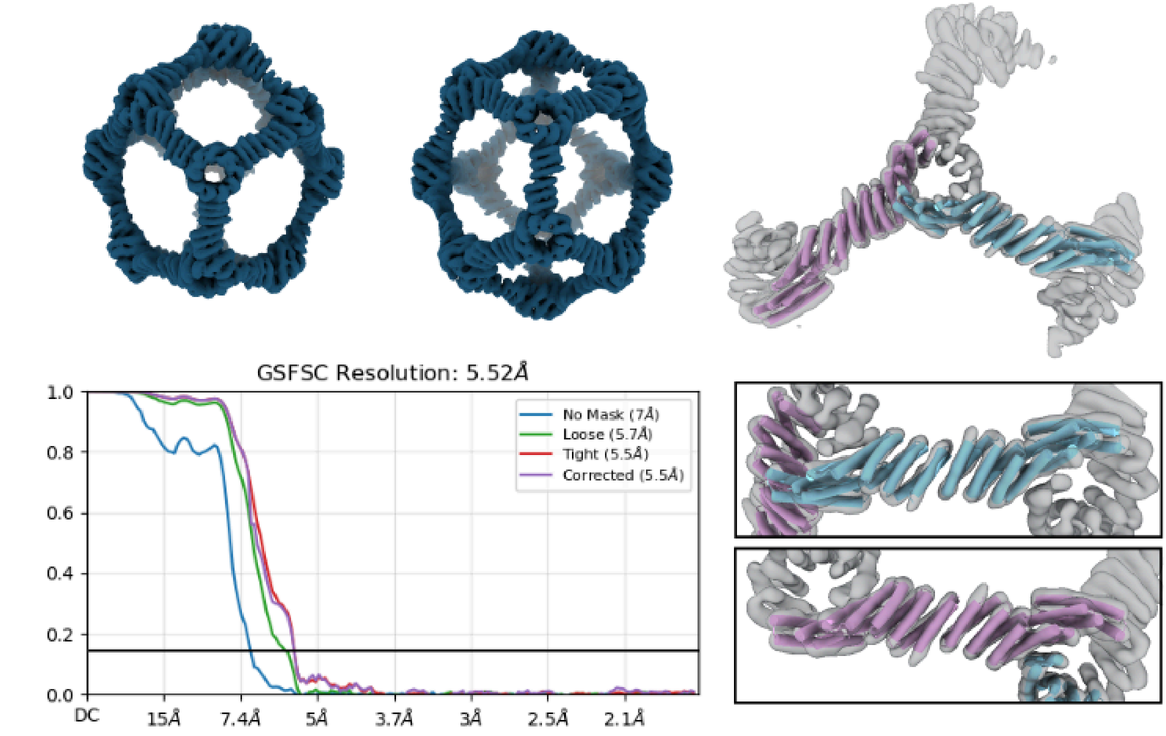


Figure S15: CryoEM processing of SL112-D D3 species. Particles were heterogeneously classified into several symmetric species as shown in Figure S8 and subsequently refined. Particles were then symmetry expanded and refined into a local model with both 4-5 and 5-5 edged of the D3 cage at 5.5 Å resolution with C1 symmetry applied. Protomers of the SL112 parent design model were then fit into the density of the two edges using ChimeraX and used as design models for D3 cages.

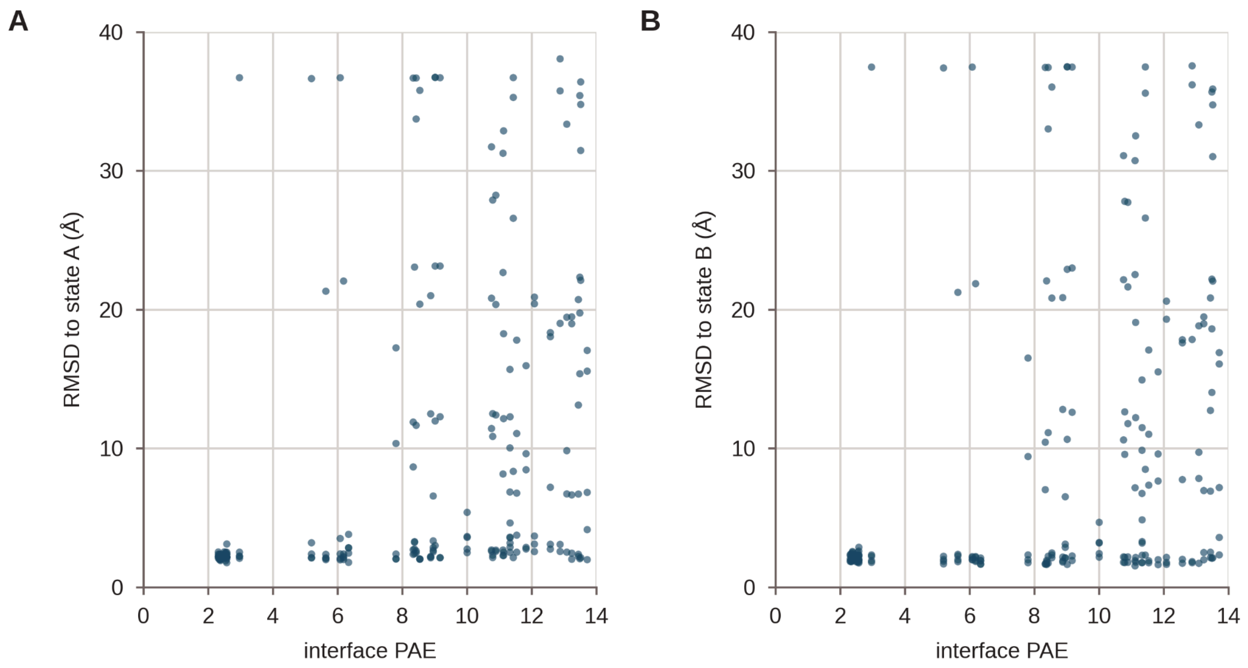


Figure S16: AF2 metrics of D3 multistate designs. Sequences designed with tied ProteinMPNN were each predicted 5 times with AF2. RMSD to the state A and state B models and pae_interface were calculated for each prediction. Predictions with RMSD less than 3 Å to both design models were selected for experimental characterization.

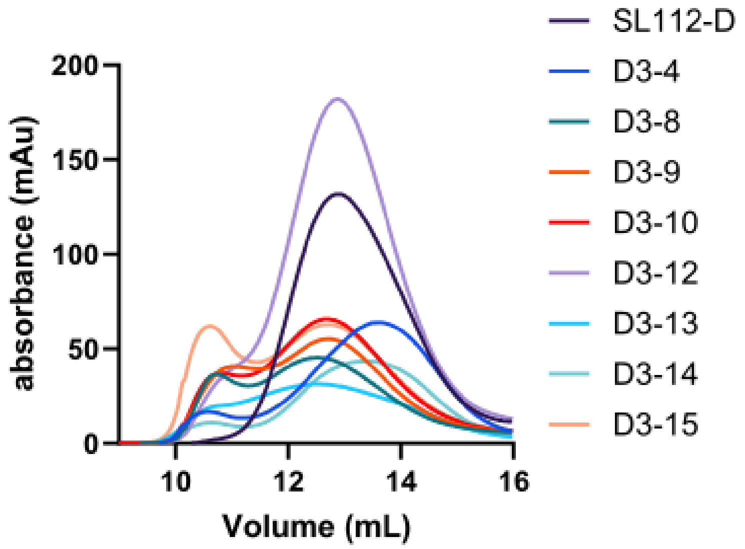


Figure S17: SEC of successful D3 designs. Purified D3 designs were analyzed via SEC and several designs eluted at the same retention time as the parent SL112-D design.

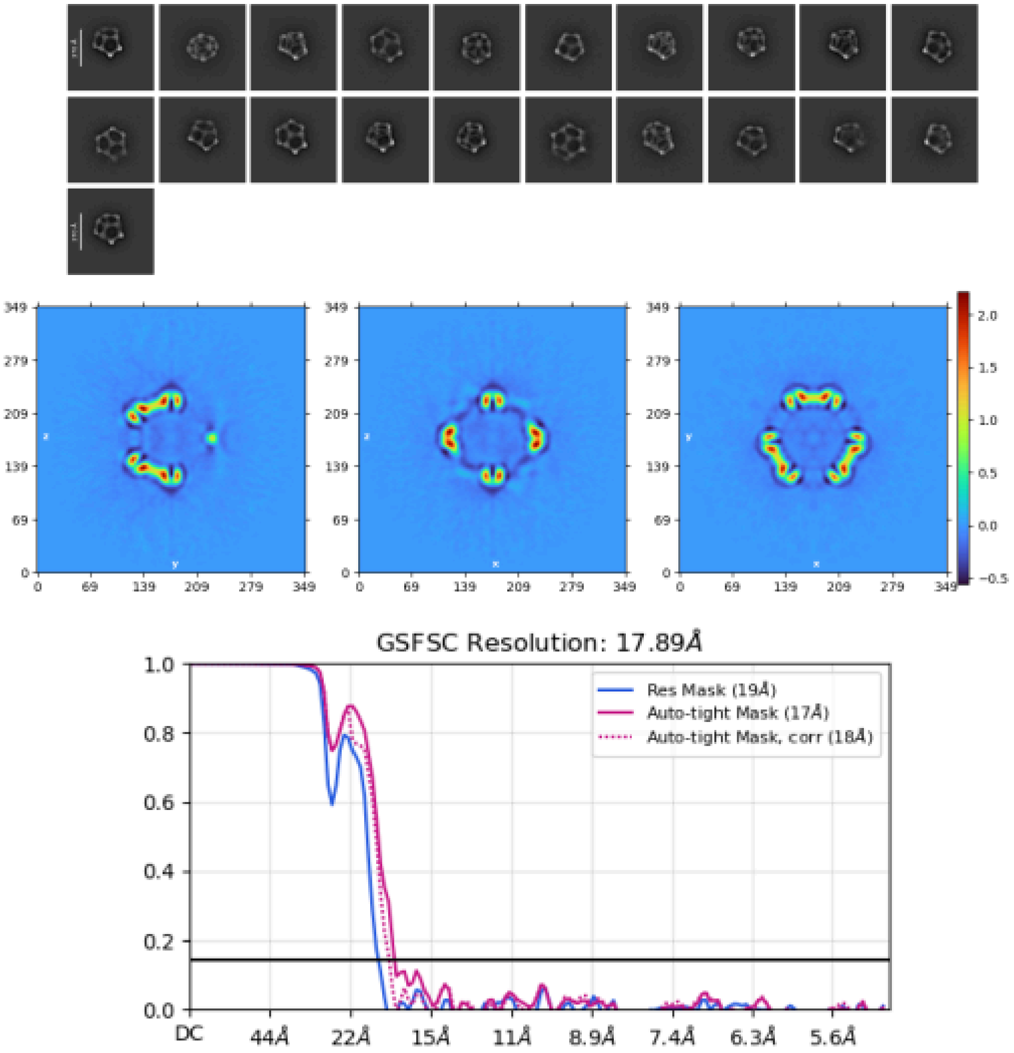


Figure S18: nsEM processing of D3-12. Particles were isolated with 2D classification and used to construct an ab initio model which was homogeneously refined with D3 symmetry.

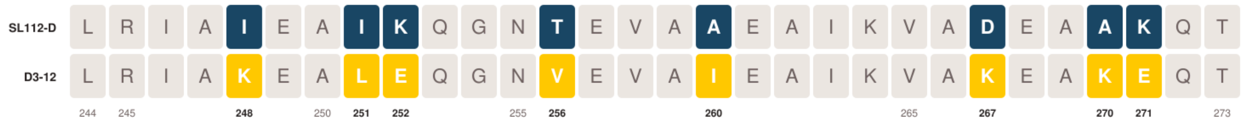


Figure S19: Sequence alignment of SL112-D and D3-12. C-terminal residues of SL112-D and D3-12 were aligned to show mutations at designed positions. Positions with a number below the residue were designed by ProteinMPNN and substitutions are colored.

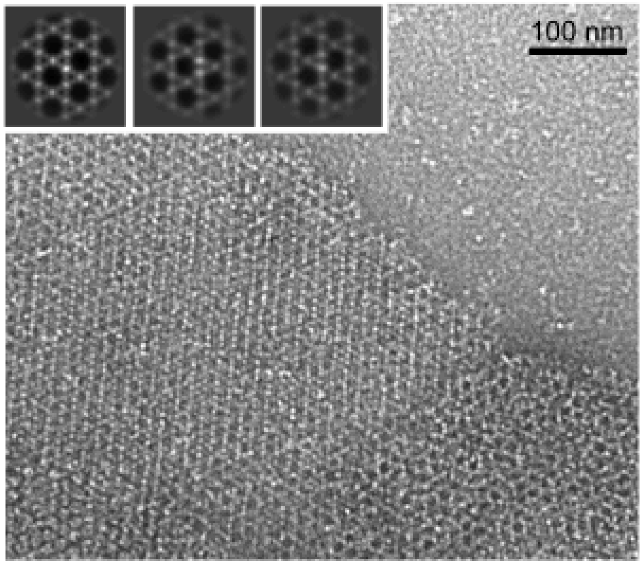


Figure S20: 2D array structure P6-01. P6-01 protomers were mixed and assembled to yield 2D arrays

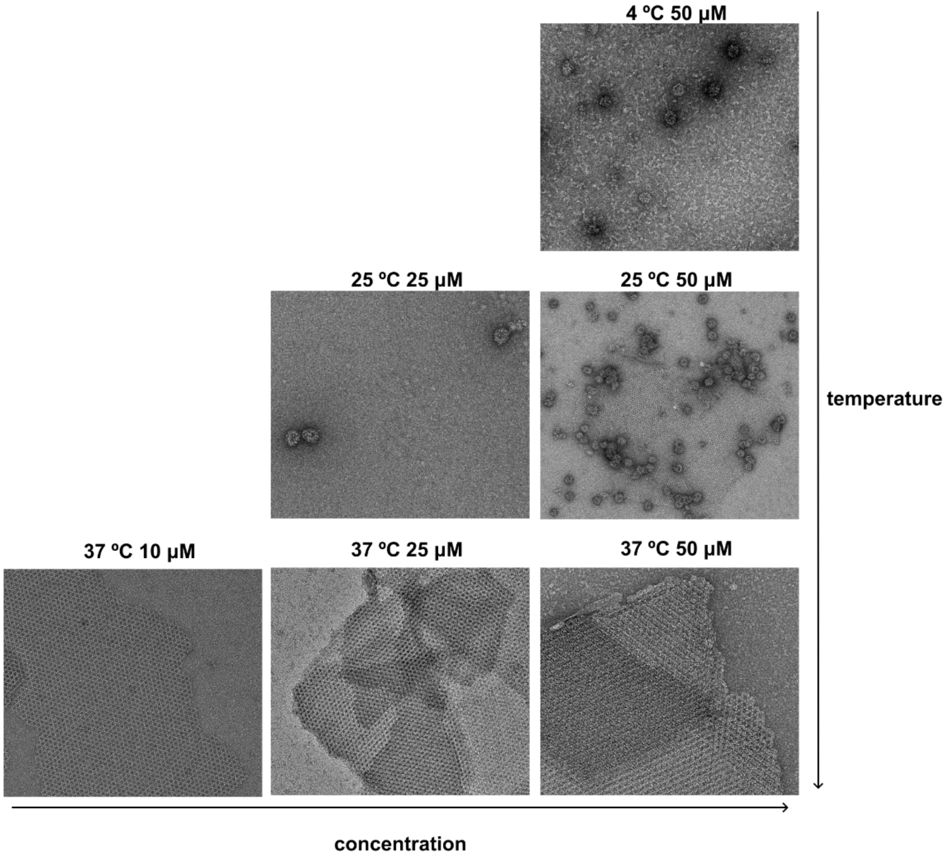


Figure S21: multi-icos 110 assembly morphology varies with assembly conditions

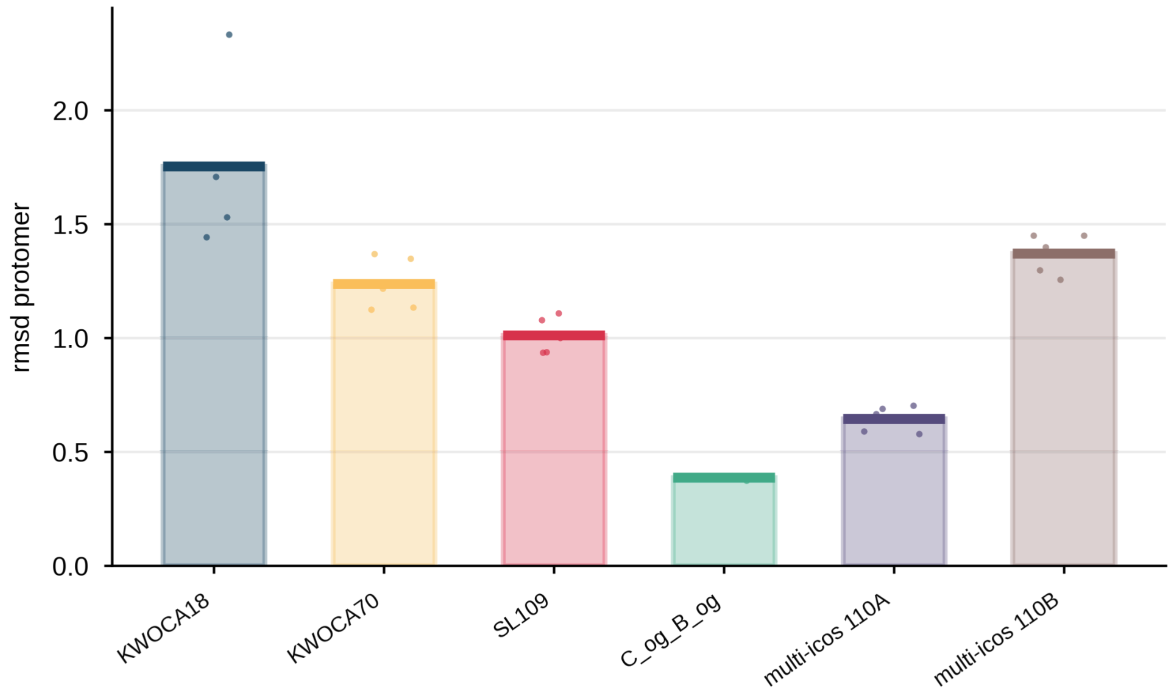


Figure S22: AF2 prediction of previously reported quasisymmetric assemblies. Designed assemblies were predicted as a 2 chain complex and the RMSD of each protomer to the protomers of the design model is reported above.
